# Harmonized single-cell landscape, intercellular crosstalk and tumor architecture of glioblastoma

**DOI:** 10.1101/2022.08.27.505439

**Authors:** Cristian Ruiz-Moreno, Sergio Marco Salas, Erik Samuelsson, Sebastian Brandner, Mariette E.G. Kranendonk, Mats Nilsson, Hendrik G. Stunnenberg

**Affiliations:** Prinses Máxima Centrum for Pediatric Oncology, Utrecht, The Netherlands; Department of Molecular Biology, Faculty of Science, Radboud University, Nijmegen, The Netherlands; Science for Life Laboratory, Department of Biochemistry and Biophysics, Stockholm University Solna, Sweden; Division of Neuropathology and Department of Neurodegenerative Disease, UCL Queen Square Institute of Neurology, Queen Square, London WC1N 3BG, United Kingdom

**Keywords:** Glioblastoma, single-cell RNA sequencing, reference mapping, transfer learning, spatial transcriptomics, *in situ* sequencing, tumor organization

## Abstract

Glioblastoma, isocitrate dehydrogenase (IDH)-wildtype (hereafter, GB), is an aggressive brain malignancy associated with a dismal prognosis and poor quality of life. Single-cell RNA sequencing has helped to grasp the complexity of the cell states and dynamic changes in GB. Large-scale data integration can help to uncover unexplored tumor pathobiology. Here, we resolved the composition of the tumor milieu and created a cellular map of GB (‘GBmap’), a curated resource that harmonizes 26 datasets gathering 240 patients and spanning over 1.1 million cells. We showcase the applications of our resource for reference mapping, transfer learning, and biological discoveries. Our results uncover the sources of pro-angiogenic signaling and the multifaceted role of mesenchymal-like cancer cells. Reconstructing the tumor architecture using spatially resolved transcriptomics unveiled a high level of well-structured neoplastic niches. The GBmap represents a framework that allows the streamlined integration and interpretation of new data and provides a platform for exploratory analysis, hypothesis generation and testing.

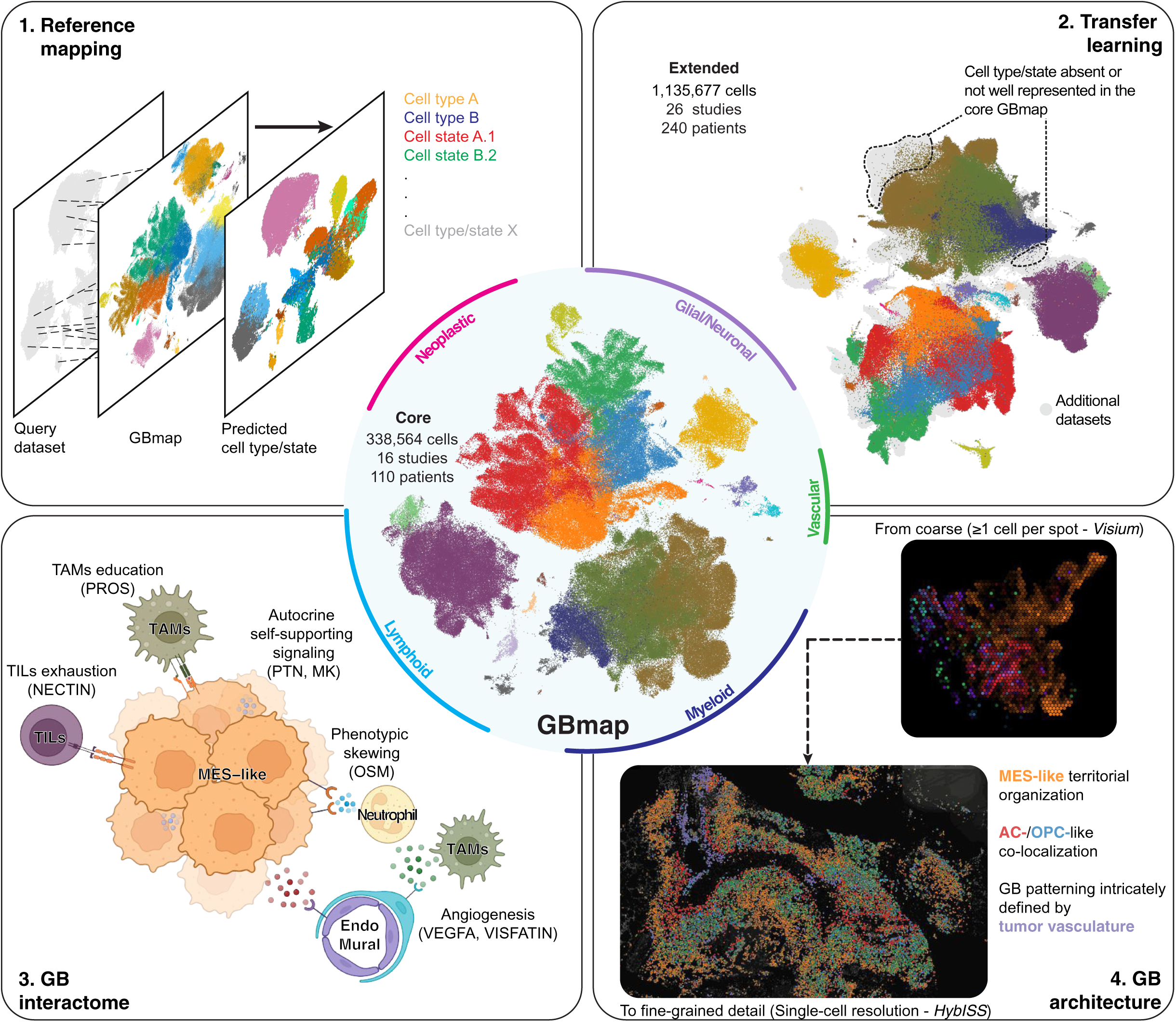

## INTRODUCTION

GB is the most common malignant brain neoplasm in adults and continues to have a poor prognosis despite developments in multimodal therapy (Miller et al., 2021). Recent advances in the molecular profiling of individual cells have unraveled a remarkable cell heterogeneity of the neoplastic cells and the tumor microenvironment (TME), especially the immune compartment (Andersen et al., 2021; Cordell et al., 2022; Suva and Tirosh, 2020). Our current understanding of the complexity, particularly at the gene expression level, points toward a dynamic process where cells transit among different states in a continuous spectrum rather than fitting unique delineated categories (Couturier et al., 2020; Garofano et al., 2021; Johnson et al., 2021; Neftel et al., 2019; Richards et al., 2021).

To better capture the molecular underpinnings of transcriptomic variation between different tumor samples, it is necessary to gather and integrate large cohorts that can empower understanding of the disease mechanisms and capture the phenotypic features of GB. Single-cell RNA sequencing (scRNA-seq) has been the leading method to characterize the tumor cellular makeup, and its adoption to study gliomas has led to the generation of a plethora of datasets that vary in size, sequencing protocols, cell enrichment, and regional sampling, amongst others (Hernandez Martinez et al., 2022). Successful harmonization of such diverse datasets can help to chart with precision the TME at single-cell resolution. Integrating scRNA-seq data has been one of the main challenges in the field, and reducing the technical variance is paramount in generating a high-quality reference map (Argelaguet et al., 2021; Luecken et al., 2022). Recent developments in computational methods have addressed these issues and allowed the generation of large-scale reference atlases in health and disease of several tissues and organs (Emont et al., 2022; Litviňuková et al., 2020; Sikkema et al., 2022).

Various studies have featured the potential of scRNA-seq to model intercellular communication by measuring the expression of ligands and receptors (L-R) in multiple cell types and disentangle communication networks (Efremova et al., 2020; Jin et al., 2021). Attempts have been made to model the cell-cell interactions in GB (Caruso et al., 2020; Hara et al., 2021; Xiao et al., 2022; Yu et al., 2020). However, they were either limited to small sample size or did not provide a holistic overview of the GB cellular composition to construct the tumor interactome. Additionally, despite current technological advances, transcriptomic profiling in a spatially resolved manner has only lately been used to explore GB (Ravi et al., 2021; Ravi et al., 2022).

We set out to build the ‘GBmap’ (**Fig. 1a**). At its core, it integrates multiple scRNA-seq datasets, providing a harmonized annotation at different levels. We demonstrate the applicability of the GBmap to annotate newly generated data robustly. Through transfer learning, we gathered more than 1.1 million cells, which increased the likelihood of identifying under-represented cell (sub)types. The GBmap facilitates the construction of a global network of cell-cell interactions. In addition, it assisted in the fine mapping of cell types and states in low-resolution spatially resolved transcriptomics (ST) data to chart the architecture of GB. Finally, *in situ* sequencing unveiled the territorial organization of the neoplastic cells at single-cell resolution. Delving into the disposition of the different cell (sub)types within the TME will deepen our knowledge of cellular neighboring patterns and crosstalk in proximal cell subpopulations.

**Figure 1.**
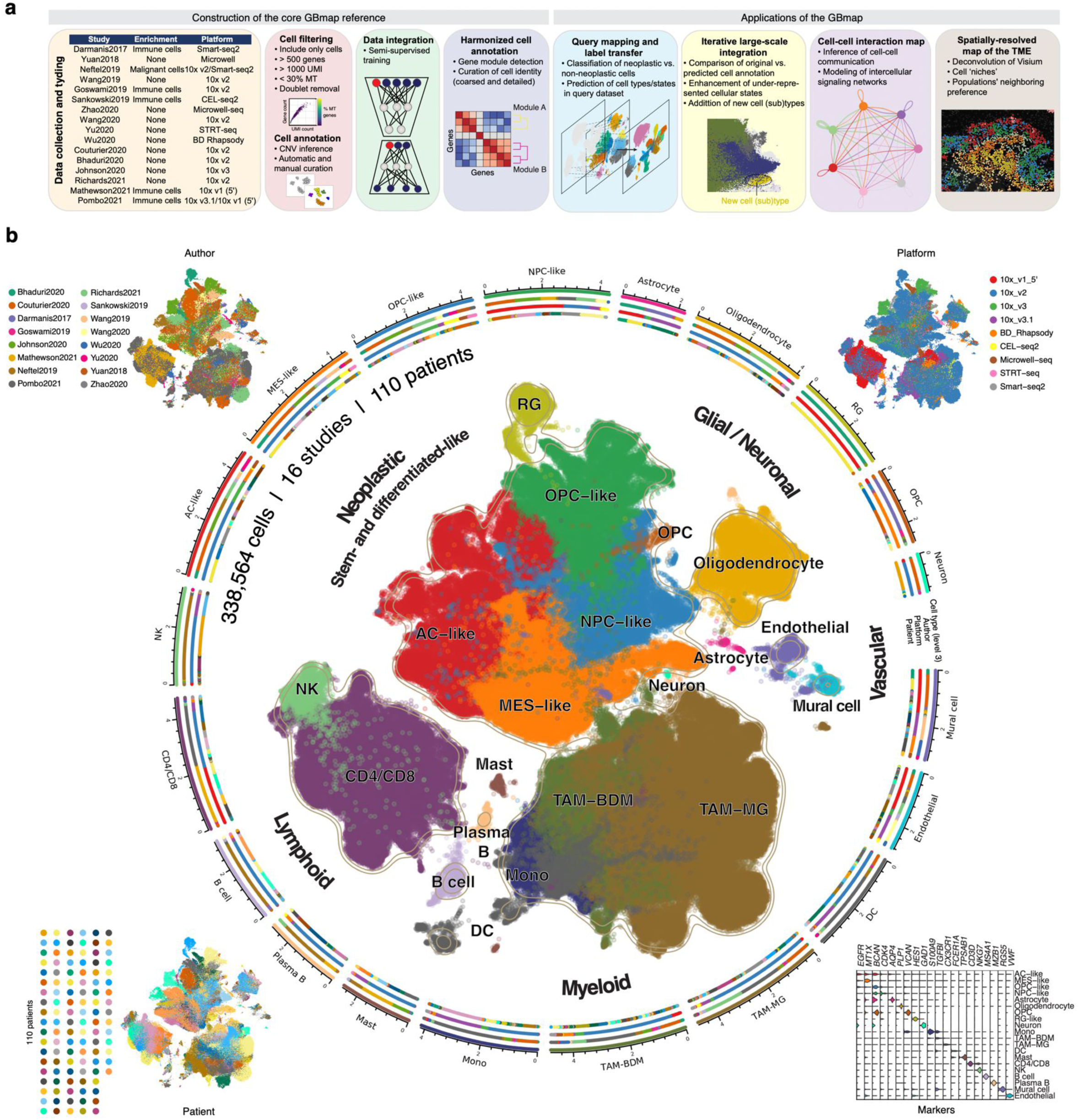
Construction of the core GBmap reference. (a) Study design and computational analysis summary. (b) UMAP representation of the GB atlas (core GBmap) over ∼330,000 cells after data integration and batch correction using SCANVI (Xu, et al., 2021). The corner insets and colored radial tracks depict the author, sequencing platform, patient, and marker-gene expression. The axis outside the circular plot shows the log scale of the total cell number for each cell type (level-3 annotation). See also S1.

## RESULTS

### Creation of a harmonized single-cell GB core reference

To establish a GB core reference, we initially gathered scRNA-seq profiles from 16 studies across multiple platforms and diverse sample preparation strategies (**Fig. 1a; Table S1**), obtaining over 330,000 cells from 110 patients. As a first step, we performed an independent automated cell annotation using a cataloged list of markers (**Table S2**), unsupervised manual assignment, and curation of broad cell types (Clarke et al., 2021). For datasets where whole-tumor profiling was performed, inference of copy number variation was assessed individually, distinguishing cells with aneuploid changes to assign non-neoplastic and neoplastic cells (**Fig. S1a**). To overcome the technical challenge of data integration, we employed a semi-supervised neural network model (scANVI) (Xu et al., 2021) implemented in the transfer-learning framework of the scArches algorithm (Lotfollahi et al., 2021), which takes advantage of the uniform prior cell type labeling to harmonize the datasets while preserving cell biodiversity. After co-embedding all cells in a dimensionality reduction space, we reconstructed a detailed TME cell map broadly divided into neoplastic and non-neoplastic cells (neuronal/glial, myeloid, lymphoid, and vascular) (**Fig. 1b**). We kept all clinical (e.g., sex, age, tumor location) and diagnostic (e.g., *EGFR* and *MGMT* status, *TP53* mutation) information whenever available (**Fig. S1a and S1b**).

To systematically segregate the major populations, we initially characterized the cell identity at a low granularity level (level-2 and -3 annotation, **Fig. 1b and S1c**). Neoplastic cells (38%), recognized by the overall presence of (inferred) copy number variation (iCNV), globally converged into two central cellular phenotypes: stem/progenitor- and differentiated-like cancer cells. The innate immune compartment (39%) could be broadly divided into dendritic cells (DC) and tumor-associated macrophages (TAMs), the latter divided by the closest transcriptomic cell ontogeny into blood-derived monocytes/macrophages (BDM) and microglia (MG) populations. Tumor infiltrative lymphocytes (TILs) (17%) were mainly constituted of T CD4+/CD8+ and natural killer (NK) cells, and to a lesser extent, B/plasma and mast cells. Non-neoplastic neuronal/glial (5%) and vascular cells (1%) embedded in the tumor were assigned based on their distinctive transcriptomic profile and the lack of iCNV (**Fig. 1b and S1a**).

Next, we identified gene modules (programs) to define representative phenotypes for each category in greater detail (**Table S3**). The gene modules detected in neoplastic cells were consistent with cellular states that mimic astrocyte (AC)-like, neural precursor cell (NPC)-like, oligodendrocyte percussor cell (OPC)-like, and mesenchymal (MES)-like states (Neftel et al., 2019) (**Fig. 2a and S2a)**. Additional cellular (sub)programs within the cancer phenotypes included hypoxia and major histocompatibility complex class (MHC)-II/cytokine modules, particularly enriched in MES-like cells as described in published studies (Hara et al., 2021; Neftel et al., 2019) (**Fig. S2b**). Enhanced cell proliferation was primarily seen with the OPC/NPC- and AC-like cells (**Fig. S2b**).

**Figure 2.**
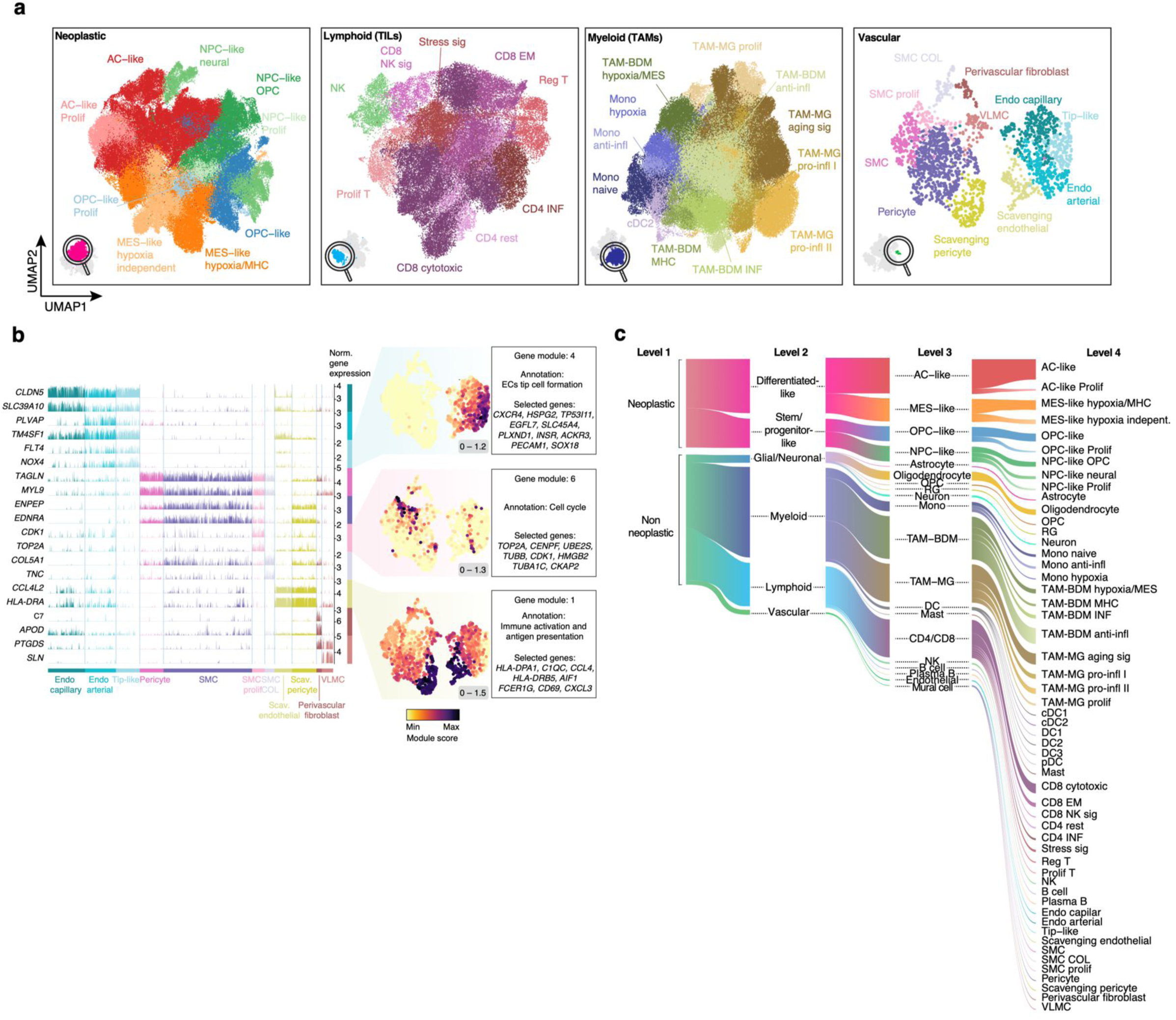
Categorization of cell states in the GBmap. (a) UMAP and cell annotation of sub-clustered neoplastic, myeloid, lymphoid, and vascular territories (level-4 annotation, detailed cell states). (b) Expression of representative genes (left) of the different cell (sub)types part of the tumor vasculature. On the right-hand side, enrichment scores of selected gene modules show signatures associated with tip-like cell formation (module 4), cell cycle (module 6), and immune activation (module 1). The minimum and maximum module scores are shown in the bottom right of each panel. (c) Overview of the GBmap cell type/state composition from coarse (level 1) to fine (level 4). See also S2.

Within the TAMs, the BDM-enriched ontogeny displayed up-regulation of gene programs associated with classical monocytes (Pombo Antunes et al., 2021), interferon (INF)-induced genes, pro-inflammatory cytokines, and tumor-supportive chemokines (**Fig. 2a and S2c**). A BDM subset characterized by the inclusion of MES-like genes (e.g., *VIM, CSTB*) matched the recently recognized MES-like myeloid phenotype (Hara et al., 2021) (**Fig. S2c and d**). Additional gene modules, such as a hypoxia-responsive program, were also enriched in the MES-like BDM subtype (**Fig. S2c**). Cells with higher expression of canonical microglial genes were divided into three principal categories: MG-like phenotypical signature with up- regulation of pro-inflammatory genes, INF response and immune activation program, and lastly, aging-like transcriptional pattern with high expression of *SPP1*, *APOE/C,* and *BIN1* (Sankowski et al., 2019) (**Fig. 2a and S2c,d**). Exploration of the lymphoid territory revealed CD4+ T cells falling into the category of regulatory (Tregs), INF signature, and effector memory cells in a polarization-state (Poch et al., 2021) (**Fig. 2a and S2e**). CD8+ T cells expressed high levels of genes associated with cytotoxicity, effector memory-associated programs, and cell proliferation. In addition, we also detected the recently discovered CD8+ NK-associated signature (Mathewson et al., 2021) defined by enrichment of *FCGR3A, GZMB,* and *KLRB1* (**Fig. S2e and f**). A stress signature enriched in heat-shock protein (HSP) genes was discernable, previously described as a genuine T cell phenotype in glioma-infiltrating T cells (Mathewson et al., 2021) (**Fig. S2e and f**).

On the vasculature, we defined cell (sub)types reported as being part of the brain blood vessels (Yang et al., 2022) as well as previously undescribed phenotypes (**Fig. 2a,b and S2g**). Endothelial cells (ECs) could be classified into tip-, capillary- and arterial-like cells based on the expression of classical zonation markers. Our large-scale analysis accurately determined that *PLVAP*, a marker consistent with fenestrated morphology typically seen in high-permeable capillaries and venous vessels, was upregulated by arterial-like GB ECs (**Fig. 2b**). *PLVAP*^hi^ arterial-like ECs suggest a novel pathological subtype in brain tumors. Mural cells consisted of smooth muscle cells (SMCs), pericytes, perivascular fibroblasts, and vascular leptomeningeal cells (**Fig. 2b and S2g,h**). Upregulation of vascular membrane remodeling (collagen and metalloproteinases), pro-angiogenic genes (e.g., *TNC*) (Rupp et al., 2016), and proliferative markers were associated with two different subpopulations of SMCs. These phenotypes have not been reported and are consistent with the enhanced microvascular proliferation in GB (**Fig. 2b**). We also identified an unconventional phenotype featured by immune activation and antigen presentation (**Fig. 2b**). The presence of an immune-enriched signature was recently discovered in a single-cell analysis of ECs in GB (Xie et al., 2021). The integrated GBmap helped to recognize a subcluster of pericytes that also has a so-called scavenging signature, which has not been described elsewhere. As suggested in other tumor types, these scavenging cells might be induced by tumor-secreted cytokines (Goveia et al., 2020).

In summary, we created GBmap, a robust core reference atlas that delineates cell types and states. We also provide a systematic annotation at different levels of granularity, from coarse to refined cell identity (**Fig. 2c and Table S4**).

### Reference mapping and large-scale integration by transfer learning

One of the main goals of building a detailed GBmap is to provide a resource for the community to project unannotated ‘query’ datasets onto a harmonized reference. To test the ability of our GBmap to recognize each cell (sub)type accurately, we generated single-nuclei RNA-seq from 11 GB cases (**Fig. 3a**, **Table S5**), obtaining 39,355 high-quality cells. We compared the manual annotation of our *de novo* dataset with the label transfer upon reference mapping (**Fig. 3b and S3a,b)**. The GBmap not only allowed the unsupervised reiteration of the major cell types but also resolved the difference between non-neoplastic and neoplastic cell types in one step (presence/absence of iCNV) (**Fig. S3c**). This is particularly important when distinguishing between states that share common gene signatures (e.g., normal OPC vs. OPC-like malignant).

**Figure 3.**
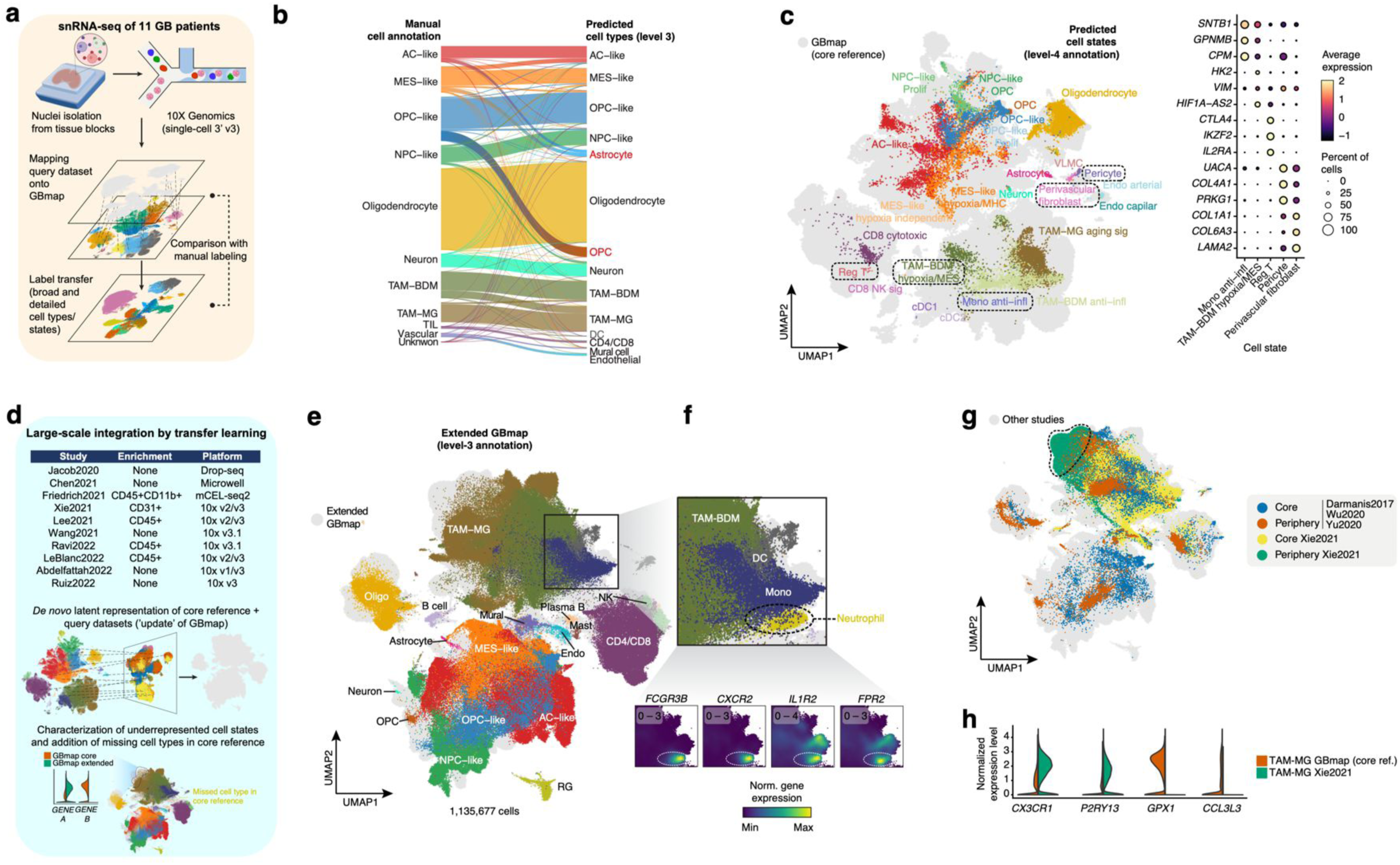
Mapping and transfer learning capabilities of the GBmap. (a) Schematic outlines the experimental and *in silico* pipeline. (b) Sankey plot comparing manual cell labeling and predicted cell types using the GBmap from 11 de novo GB patients (this study) using an anchoring method for reference mapping (Azimuth) (Hao, et al., 2021). (c) Projection and cell type prediction of detailed cell states of *de novo* mapped query cells (this study) onto the GBmap UMAP. On the right, expression of gene markers from selected cell states. (d) Diagram of comparison and machine-learning process applied to public and our own dataset. (e) Updated UMAP of the integrated GBmap and query datasets (extended GBmap). Cells from the integrated studies are colored gray using the neural network model developed in scArches (Lotfollahi, et al., 2021). Cells from the GBmap core reference are colored by level-3 annotation. (f) Close-up of UMAP region identifying a neutrophil cluster not present in the core GBmap. Below is the expression of classical neutrophil gene markers. The minimum and maximum normalized gene expression values are shown in the top left of each panel. (g) Projection of multi-sector biopsy datasets onto the extended GBmap, colored by region (core or periphery). (h) Expression of selected phenotypical microglia (CX3CR1, P2RY13) and tumor-induced genes (GPX1, CCL3L3) contrasting TAM-MG from the core GBmap and the predicted microglial cells on the Xie2021 dataset. See also S3.

The GBmap also transferred the information of fine cell subtypes to uncover phenotypes that otherwise would have remained hidden. We could identify TIL subsets in our 11 GB sample set with an expression pattern of regulatory T cells, typified by the high expression of *CTLA4* and *IKZF2* (**Fig. 3c**). This population was not detected without the label transfer provided by the GBmap, not even by sub-setting and re-clustering (**Fig. S3d**). Likewise, other cell subtypes that could not be confidently classified in the standalone analysis of our 11-sample cohort could now be correctly recognized, including anti-inflammatory monocytes, hypoxia/MES-like TAM-BDM, and perivascular fibroblasts (**Fig. 3c**). Taken together, the GBmap reference facilitated robust identification of the cellular composition in newly generated data.

It is anticipated that the GBmap will ‘grow’ over time as new studies will capture cell (sub)types that are not well or at all represented in the core GBmap reference. Projecting new datasets on a predetermined reference map, although practical for fast evaluation and interpretation, has the disadvantage of forcing the queried cells to the reference and hiding novel findings. The transfer-learning method developed in the scArches pipeline instead updates and extends a trained model overcoming the limitation of forced cell mapping. To exemplify the integration of newly generated single-cell datasets into the core GBmap, we ‘upgraded’ it with new datasets that became available after the construction of our trained model, including our own profiled GB tissues (**Table S1**) (**Fig. 3d**). The outcome generated a joint embedding resulting in the extended GBmap (>1.1 million cells) (**Fig. 3e**). Our model recapitulated the major cell groups when comparing the label transfer from the core reference (predicted cell type) to the newly mapped datasets (original annotation) enabling consensual annotation across studies **(Fig. S3f**). Of note, after large-scale integration, we detected a cluster with a high expression of neutrophil markers (e.g., *FCGR3B, CXCR2, FPR2*) (**Fig. 3f**). Importantly, tumor-associated neutrophils were absent in the core GBmap reference and could now be retrieved in the queried datasets.

Comparing the TAM-MG cells in the core GBmap (**Fig. 3g**), we found a subset of MG cells in the extended GBmap that expressed higher levels of bona fide markers of a core transcriptional microglial signature (e.g., *CX3CR1*, *P2RY13*) and lower expression of catabolic (e.g., *GPX1*) or cell activation (e.g., *CCL3L3*) processes (**Fig. 3h**). These cells mainly belonged to the integrated Xie2021 dataset (**Fig. 3g and S3g**). The multi-sector biopsy performed by Xie *et al*. showed that the histological assessment of the neighboring tumor tissue largely concurred with normal brain microanatomy. In our analysis of the extended GBmap, we could determine that in the peripheral area, their data was enriched for homeostatic resident MG cells. Due to the higher number of cells profiled by Xie *et al*. compared to previous multi-sector biopsy studies included in the core reference (Darmanis et al., 2017; Wu et al., 2020; Yu et al., 2020) (**Fig. S3h**), the divergence between ‘naïve’ resident microglia and TAM-MG was drawn out during the transfer learning process.

These results show the utility of the GBmap in adding depth to hidden or overlooked findings of cell phenotypes. It provides a framework that enables the discovery of (disease) specific cell states and can ‘learn’ when new data is added. Of note, all subsequent analyses were carried out on the extended GBmap.

### GB interactome uncovers signaling implicated in phenotypic turnover, immunomodulation, and angiogenesis

The crosstalk between neoplastic and non-neoplastic cells in the TME is thought to be pivotal in preserving cellular plasticity and impacting tumor growth and progression. We used the extended GBmap to build a comprehensive cell-cell communication network based on the weighted expression of L-R pairs in each cell type. To identify which interaction pathways are likely to change upon neoplastic transformation, we compared the inferred *in silico* crosstalk of the extended GBmap with a curated spatiotemporal cell atlas of the human brain (Song et al., 2021) (**Fig. S4a)**.

Following L-R pairs inference in each condition, we analyzed them together via joint manifold learning and performed a classification of the detected networks based on their communication similarity. In total, 110 pathways encompassing 1041 significant L-R pairs were detected, sorted into six main groups, and projected onto a shared two-dimensional space (**Fig. 4a**). Four of the six groups shared pathways between healthy and tumor. GB-only communication pathways uniquely formed groups 2 and 4. The spectrum of signaling networks linked to GB involved immune chemoattraction, vascular proliferation, and tumor maintenance (**Fig. 4a; S4b**). By computing the Euclidean distance between the shared signaling patterns (58/110 pathways), we observed a considerable distance for pathways that, under non-pathological conditions, support cell functions of resident brain cells such as NOTCH (Ables et al., 2011), junctional adhesion molecules (JAM) (Ebnet, 2017), and apolipoprotein E (APOE) (Flowers and Rebeck, 2020) (**Fig. 4b**). Increased pathway distances point toward changes in how sender and receiver cells handle a given signaling pathway. This might suggest that cancer cells hijack and reprogram homeostatic cellular brain processes to boost tumor growth and development.

**Figure 4.**
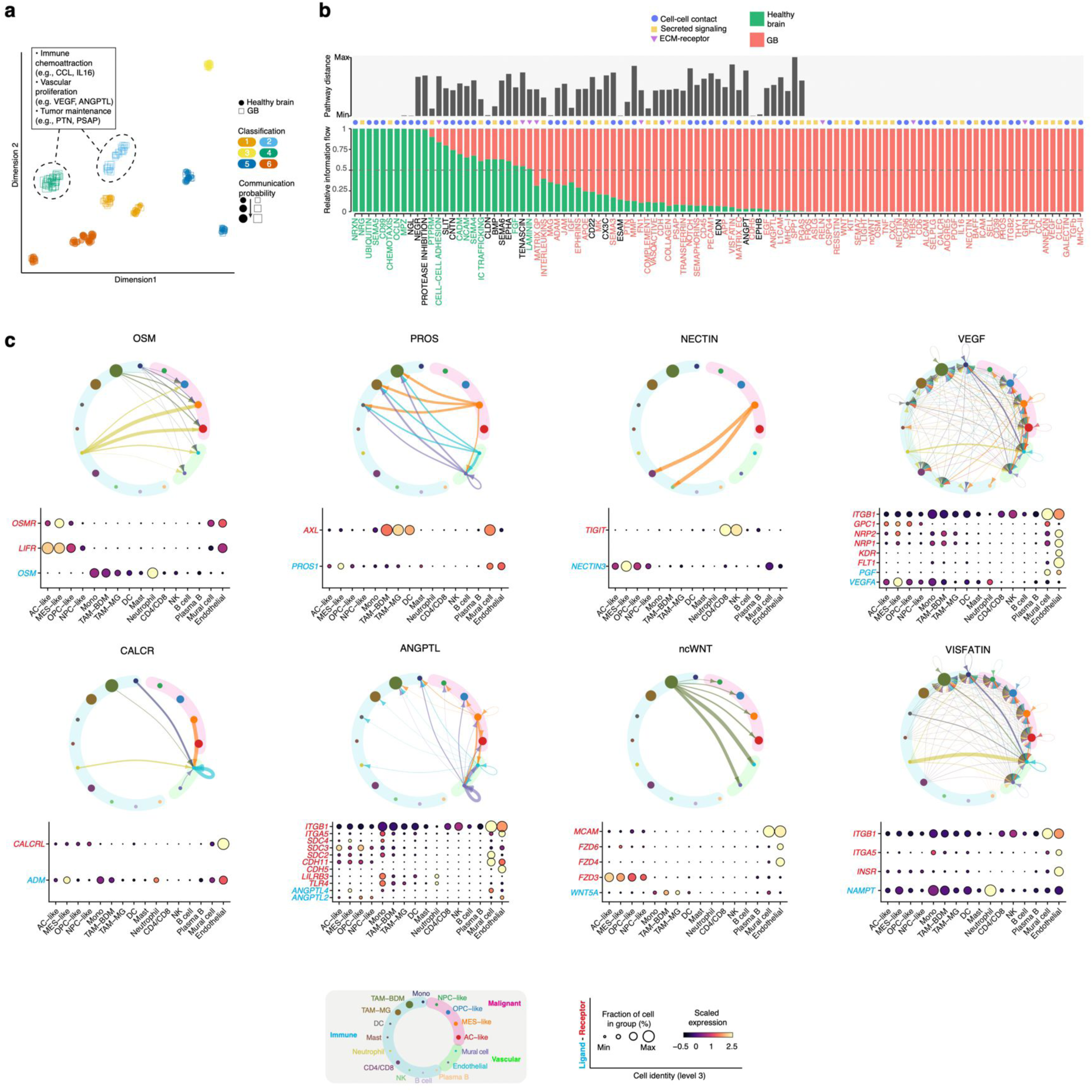
GB interactome. (a) Shared two-dimensional manifold of jointly projected and clustered signaling pathways from healthy brain and GB according to their structural similarity. (b) Bar plot of the significant signaling pathways ranked based on their differences in overall information flow within the inferred networks between healthy brain (Song, et al., 2021) and GB. Pairwise Euclidean distance in the shared two-dimensional manifold and pathway category are shown above for each communication network. The signaling pathways colored green are more enriched in the healthy brain, the ones colored black are equally enriched in the healthy brain and GB, and the ones colored red are more enriched in GB. (c) Circos plots depict the inferred signaling network interaction among neoplastic, immune, and vascular cells. Circle sizes are proportional to the number of cells in each cell cluster (level-3 annotation) on the extended GBmap, and edge width represents the communication probability. On the bottom, scaled gene expression of ligand (blue) and receptor (red) pairs of each selected communication pathway. See also S4.

Our interactome robustly recovered well-established GB-specific signaling patterns, including signaling networks among cancer cells associated with tumor growth, adaptive mechanisms of resistance, and glioma stem cell maintenance (Han et al., 2019; López-Valero et al., 2020; Osuka et al., 2021; Shi et al., 2017) (EGF, PTN, CDH, MK). Among immune cells, the inferred L-R pairs belong primarily to chemokine/cytokine families (e.g., CXCR-CXCXL families), complement system, antigen presentation (e.g., MHC genes), and adhesion molecules (e.g., ICAM and ALCAM) (**Fig. 4b and S4c**). In addition, our L-R predictions hint at novel players, such as neutrophils, that could mold the TME and communication networks covering distinct biological and histological features (e.g., microvascular proliferation). A recent study illustrated that the (mal)functional interaction between TAMs and cancer cells through the oncostatin M (OSM) pathway could shape the malignant phenotype towards an MES-like state (Hara et al., 2021). We found that interactions involved in the OSM pathway were not limited to TAMs. To a large extent, neutrophils express OSM (**Fig. 4c**), which is further up-regulated in the presence of TAMs (Zhou et al., 2021). Neutrophils account for up to 20% of the immune cells in GB (Klemm et al., 2020) and hence could potentially impact cellular communication within the tumor (**Fig. 4c and S4c**).

The inferred interplay between MES-like neoplastic cells (and, to a lesser extent, vascular cells) with TAMs via PROS pathway (PROS1-AXL) suggests a potential mechanism by which tumor cells may inhibit pro-inflammatory TAMs polarization (Ubil et al., 2018) (**Fig. 4c**). Predominant expression of *NECTIN3* by the MES-like cells can sustain an exhausted T-cell phenotype upon binding to *TIGIT* (Reches et al., 2020) (**Fig. 4c**), an immunoglobulin inhibitory receptor expressed in TILs. In line with a recent investigation (Xiao et al., 2022), MES-like cancer cells are dominant mediators of signaling growth factors that stimulate neovascularization (VEGF, ANGPTL, CALCR), with more discrete participation of monocyte-/TAM-BDM-hypoxia-associated cells (**Fig. 4c; S4c**). Their pro-angiogenic role is likely a feedforward mechanism to hypoxic metabolic cues (**Fig. S4d and S4e)**.

Our *in silico* communication modeling hints at additional pro-angiogenic signaling influx through non-conventional pathways. TAM-BDM selectively expresses wingless-related integration site family member 5A (*WNT5A*), which upon interaction with Frizzled family receptors and CD146 (*MCAM*), can trigger new blood vessel sprouting (Chen et al., 2021) (**Fig. 4c**). In addition, we found that neutrophils are the primary source of *NAMPT*, part of the VISFATIN pathway (**Fig. 4c**), which is critical for the pro-angiogenic activity in other solid tumors (Pylaeva et al., 2019) and could also be an essential vascular mediator in GB.

Overall, the GBmap provided the structural basis to create the signaling network within the TME. The analysis provides support for signaling events of well-established as well as new roles in GB. We highlight the inferred influence of MES-like cancer cells in immunomodulation and neovascularization. We find that the roots of pro-angiogenic crosstalk in the TME are not limited to MES-like neoplastic cells but are also reinforced by TAMs/neutrophils.

### Regional organization of the GB TME

Tissue organization and architecture influence cell behavior, and cell-cell interactions can impact tumor pathogenesis. Our findings on the communication network highlight that properties of the GB micro-ecosystem, such as restricted oxygen access, can trigger signaling in response to environmental cues. This likely reflects how cells are arranged within the tumor and who their neighbors are. Recent spatial scrutiny of GB tissues deciphered distinct transcriptional states, ranging from progenitor/development-like to reactive immune- and hypoxia-related programs (Ravi et al., 2022). Precise mapping of the cellular localization of the TME components using the GBmap could help to refine the tumor structure and delineate cellular crosstalk between tissue neighborhoods (**Fig. 5a**). Employing a Bayesian model for spatial multi-resolution mapping (Cell2location) (Kleshchevnikov et al., 2022), we determined the cell-type frequency in each capture area and charted the global architecture of GB in publicly available ST datasets (STAR Methods).

**Figure 5.**
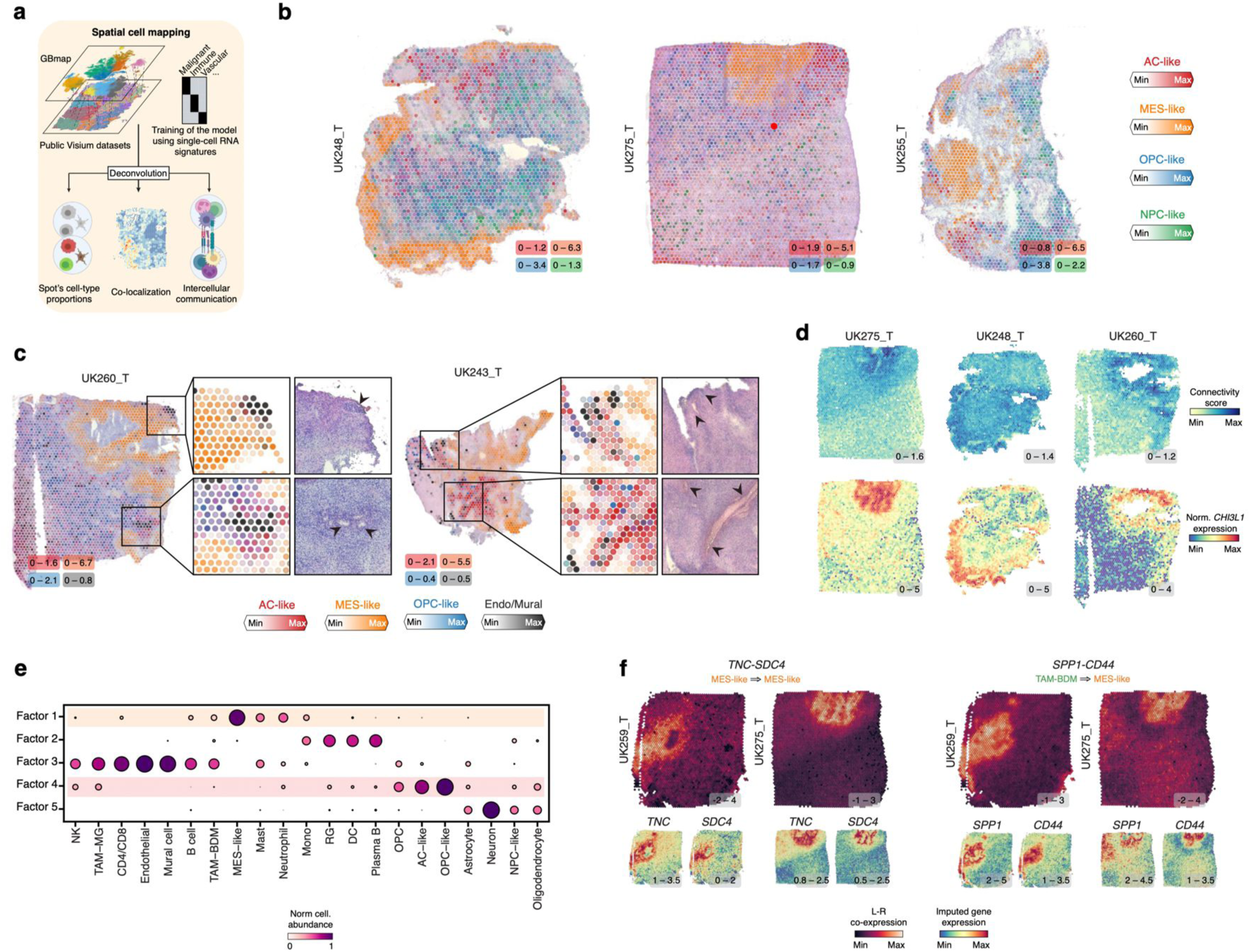
Spatial mapping and cell signaling of GB cancer cells and the TME. (a) Schematic view of the *in silico* deconvolution pipeline. (b) Estimated cell abundance (color intensity) of the four main cancer cell states is shown over the H&E image of three primary GB. The minimum and maximum cell abundances are shown in the bottom right of each panel. (c) Estimated cell abundance (color intensity) of the AC-, OPC- and MES-like cancer cell states and vascular cells are shown over the H&E image of two primary GB. Boxes highlight the areas shown in the inset panels. The left boxes show the estimated cell abundance, and the right boxes depict the H&E staining. Black arrowheads point to blood vessels embedded within the tumor. The minimum and maximum cell abundances are shown in the bottom left of each panel. (d) Spatial connectivity score and expression of the key connectivity gene *CHI3L1* of different GB samples are shown in Fig 5a. The minimum and maximum connectivity score and normalized gene expression are shown in the bottom right of each panel. (e) Dot plot of the estimated NMF weights of cell groups (columns - level-3 annotation) across NMF components (rows), which corresponds to cellular compartments. Colored by relative weights, normalized across components for each cell cluster. (f) Spatial maps of co-localized gene expression of key L-R pairs. Bottom panels show the expression of each L-R separately. The minimum and maximum L-R co-localization score and imputed gene expression are shown in the bottom right of each panel. See also S5.

The distribution of the cancer cell states revealed regionally conserved consistencies across multiple samples. AC- and NPC/OPC-like states showed extensive co-localization in several areas, whereas the MES-like phenotype formed demarcated ‘patches’ (**Fig. 5b and S5a**). MES-like areas were mainly populated by this phenotype, with fewer cells sharing the same space (low cell diversity) and a lack of co-localization with the other cancer phenotypes (**Fig, S5b**). We noticed that tumor samples containing regions with large, immersed blood vessels depicted a structured patterning. It appears that AC-/OPC-like cancer cells surround the main tumor blood vessels, followed by an adjacent neighborhood populated by MES-like phenotype (**Fig. 5c**). Furthermore, transition areas between AC/OPC- and MES-like resemble cancer cells from the MES-like hypoxia-independent phenotype, which later transit to a hypoxia-associated state (**Fig. S5c)**. Hypoxia may be a critical metabolic determinant of the distribution of cancer states within the tumor; however, the distinctive disposition of MES-like cancer cells might also reflect higher cell interconnectivity. We employed a recent transcriptomic connectivity signature established using a xenografted mouse model and validated in an independent cohort of primary human GB (Hai et al., 2021). We found a high connectivity score and upregulation of the molecular connectivity marker *CHI3L1* in regions dominated by MES-like malignant cells (**Fig. 5d).**

Considering the estimated cell-type abundance in each spot, we randomly screened all different tumor samples and regions and applied a non-negative matrix factorization (NMF) to delve into the spatial co-occurrence of cells that determine ’compartments’ within the TME. Factor 5 was composed mainly of normal glial and neuronal cells and usually found in areas adjoining non-tumoral tissue **(Fig. 5e)**. Factors 2 and 3 were enriched with different immune populations suggesting that some regions within the tumor may facilitate immune-immune crosstalk **(Fig. 5e)**. Factor 4 corroborated the AC/OPC-like co-localization, and factor 1 the zonal disposition of MES-like across GB samples (**Fig. 5e**). The later compartments suggest different associations with specific immune cells. MES-like cancer cells shared neighborhoods with monocytes/TAM-BDM and, to a lesser extent, TILs, corroborating previous reports (Hara et al., 2021; Mathewson et al., 2021) (**Fig. 5e**). Our analysis unveiled a modest enrichment of neutrophils and mast cells in MES-like dominated regions (**Fig. 5e**). A recent study reported an association between neutrophils and MES-like GB tumors in a mouse model (Magod et al., 2021). Our analysis indicates such a partnership also occurs in primary human GB. The AC-/OPC-like compartment (factor 4) was associated with a ‘colder’ immune environment with the contribution of NK and TAM-MG (**Fig. 5e**).

Investigating the spatial communication between neighborhoods, we found that several inferred L-R pairs were driven by MES-like dominated areas, in line with our *in silico* inference of the GB interactome (**Fig. S5e**). This suggests a prominent role of MES-like regions in supporting pro-angiogenic signaling (**Fig. S5e**). Our spatial cell-cell interaction network further indicated crosstalk of MES-like cancer cells with TAM through the osteopontin pathway (e.g., *SPP1-CD44*) as well as interactions with the extracellular matrix (ECM) via tenascin C (TNC) (**Fig. 5f**). These signaling networks could well be essential to sustain and support the well-delimited architecture displayed by the MES-like cancer phenotype (Chen et al., 2019; Sun et al., 2018; Xia et al., 2015).

Collectively, our results provide insights into the architecture of GB and illustrate global spatiotemporal patterning, with co-localization of AC- and OPC-like and a predilected self- neighboring disposition of MES-like cancer cells.

### Tumor vasculature delineates territorial allocation of cancer states

Our observations on the coarse ST data prompted us to resolve cell (co)localization at high resolution. We employed a targeted hybridization-based *in situ* sequencing (HybISS) (Gyllborg et al., 2020) that reads out the sequences of preselected transcripts within the intact tissue at single-cell resolution. We explored two GB tissues from the 11 patients profiled in our study (NH17-2680 and NH19-565) and generated padlock probes against 194 genes (**Table S6**) specific for neoplastic (four malignant states) and non-neoplastic cell types (lymphoid, myeloid, and vascular). We mapped the cell types that compose the TME and assigned transcripts to cells through a probabilistic method for cell typing by *in situ* sequencing (pciSeq) (Qian et al., 2020) (**Fig. S6a,b and c**). The tissue sections corroborated and significantly extended the phenotypic patterning uncovered by the low granularity ST data (**Fig. 6a; S6d**). Unique patches dominated by MES-like cells or mixed AC/OPC/NPC-like were seen throughout the section (**Fig. 6a, boxes I, II**). Notably, the GB cancer phenotypes showed a layered distribution relative to the endothelial and mural cells (**Fig. 6a, boxes II, III, IV**). AC-like cells were primarily enriched near the endothelium, whereas MES-like cells were often layered further away and characterized by high expression of hypoxia and pro-angiogenic genes (e.g., *VEGFA, HILPDA*) (**Fig. 6b**). The OPC/NPC layer near the blood vessels (**Fig. 6a, boxes I, II)** is in accordance with the published description of a perivascular niche that is enriched for cancer stem-like cells (Adjei-Sowah et al., 2022; Jung et al., 2021).

**Figure 6.**
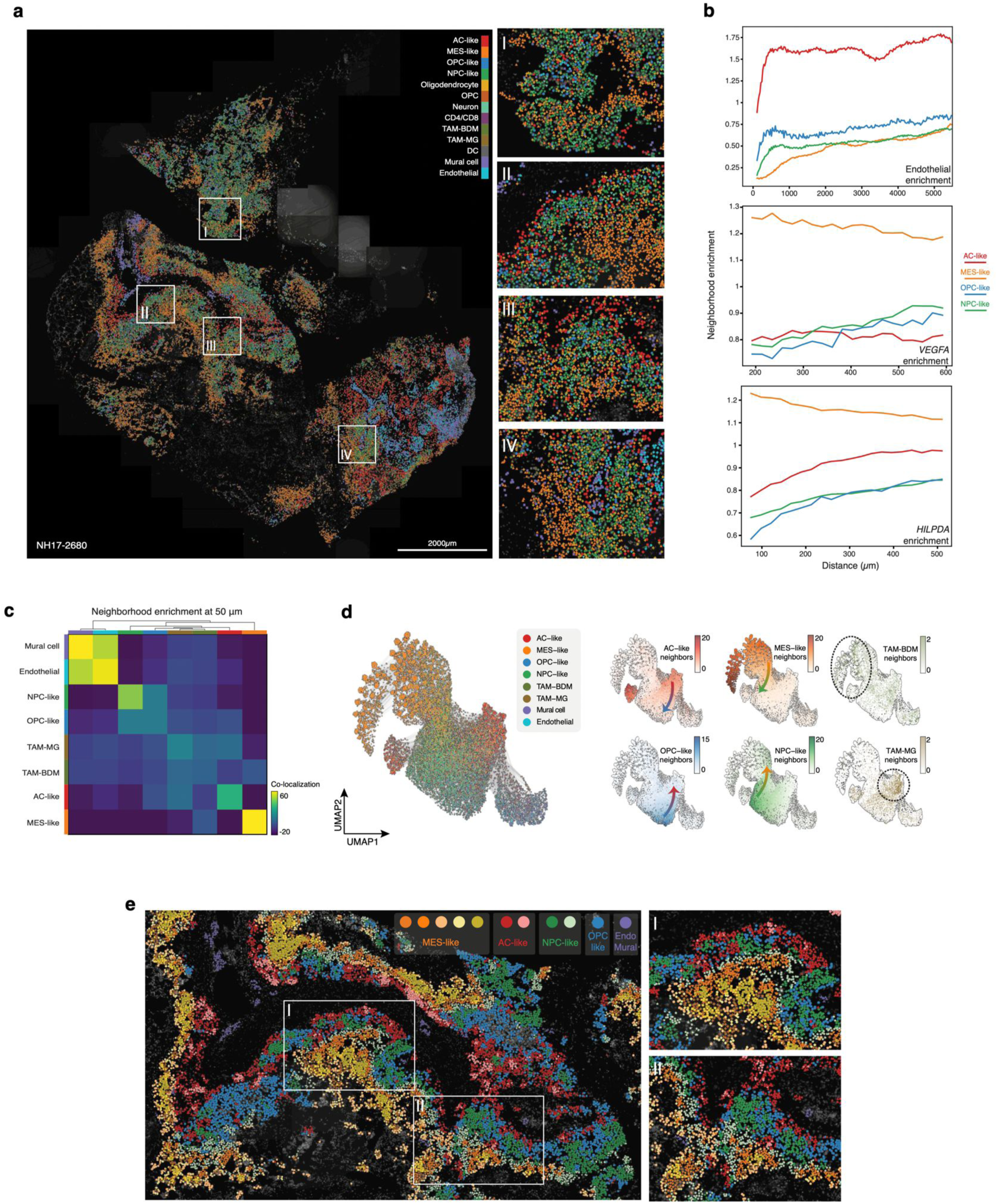
Reconstruction of the GB architecture at single-cell resolution. (a) pciSeq analysis of GB tissue (NH17-2680) showing the distribution of cell types and states, colored according to cell identity. Boxes I-IV shows regions of interest, identifying areas of specific cellular architecture (I-IV). (b) Distribution of distances (x-axis) and neighborhood enrichment (y-axis) of malignant phenotypes to endothelial cells expressing *VEGFA*, and *HILPDA*. Neighborhood enrichment (y-axis) is the probability of finding a spatial signal for gene/cell type A when cell type B is present at X distance (x-axis). (c) Heatmap of clustering of neighborhood enrichment among cell types within a 50μm radius from each cell. (d) On the left-hand side, UMAP clustering of spatial analysis of the composition of the neighboring pattern identified within the range of 50 μm from each cell, colored by cell type. On the right, the distribution of the malignant states and TAMs across the range of cellular niches, colored by cell density. Arrows depict preferential spatial transitions among cancer states. The dotted line highlights the areas of myeloid enrichment (TAM-BDM and TAM-MG). (e) Spatial map of preferential nearest neighbors of the neoplastic cells (see Methods), colored by sub-niches. Boxes I and II show zoomed-in areas on two areas covering the range of malignant sub-niches from perivascular to areas distant from the vasculature. See also S6.

To gain further insight into the cellular patches, we performed a neighborhood enrichment analysis and clustered the cells based on their nearest neighbors within a 50 µm radius (**Fig. 6c**). The organization of the neoplastic cells appears to follow specific state-dominated domains. We observed several neighborhood patterns; firstly, HybISS revealed an extensive intra-co-localization of MES-like cells. Secondly a frequent association between MES-like and TAM-BDM as well as AC-like with TAM-MG (**Fig. 6c and d**). Both patterns align with our findings on the Visium datasets (**Fig. 5e**). Thirdly, there are shared neighborhoods between ECs with mural cells and NPC-like with OPC-like. Cluster analysis of the neighbors showed areas of gradual connectivity between OPC-like to AC-like-enriched domains and, to a lesser extent, MES-like and NPC-like-dominated domains (**Fig. 6d**). Neftel *et al*. showed that cancer cells could intricately transit among multiple cellular states (Neftel et al., 2019). Other studies performing trajectory inference proposed an inclination of progenitor-like OPC/NPC malignant cells to shift towards differentiated inflammatory-related AC/MES phenotype (Liu et al., 2020; Ravi et al., 2021; Wang et al., 2019). Our findings suggest that within the tumor, the plasticity of the cancer cells might allow specific phenotypes transitions (e.g., OPC- to AC-like rather than to the other two states), likely favored by their surrounding habitat, which may act as the driving force of cell state commitment. Performing cluster detection of the neighboring features, our spatial analysis uncovered distinct cancer state sub-niches suggesting further diversity based on their preferential nearest neighbors and presumptively additional environmental cues within the GB ecosystem (**Fig. 6e, boxes I, II and S6e**). Taken together, the disposition of neoplastic cells alongside the ECs/mural cells suggests that the blood vessels may act as one of the factors that lead to the zoned territorial organization.

We provide the first single-cell resolution map of the spatial organization of GB and offer a detailed view of the TME architecture. Our work is an essential step toward a better understanding of the relationship between cell plasticity and tumor micro-ecosystems.

## DISCUSSION

To tackle the challenge of confidently identifying and assigning cell identities, we gathered numerous published cohorts of patients and cells to capture the entire spectrum of cellular states in GB as much as possible. We created an integrated compendium of GB using scANVI (Xu et al., 2021), effectively reducing the batch effect between datasets while retaining biological information. Beyond well-recognized and recently discovered cell states, we could pinpoint scarce cell populations, such as normal neurons (0,00006498%), and novel phenotypes, such as *PLVAP*^hi^ arterial-like ECs and immune-associated scavenging pericytes. Considering previous studies and the findings of our independent analysis, we propose a harmonized annotation for the different cells in a structured and comprehensive manner (**Fig 2c and Table S4**).

We showcase how the GBmap can accurately transfer the cell’s identity to new GB datasets and uncover phenotypes that would have remained hidden without the atlas (e.g., Tregs). We could also accurately designate which cells are neoplastic in one step, including hard-to-discern types such as non-neoplastic OPC and OPC-like tumor cells. The GBmap is an ‘ever-evolving’ platform that can continuously be improved when new information is fed into it. We demonstrated that cell types (e.g., neutrophils) and under-represented phenotypic states (e.g., naïve MG) became discernible by updating the core reference with additional datasets. New studies focusing on specific cell types/states could help fill in missing pieces of the puzzle, for instance, other granulocytes or lymphatic vessel cells that could play a role in the TME.

Additional extensions of the GBmap or data exploration from a different perspective will offer complementary information to improve our understanding of tumor pathobiology.

On the analytical side, we exploited the GBmap to infer the cell-cell interaction panorama of GB. By comparing the inferred crosstalk between cells in the healthy brain and gliomas, we found that the tumor might subvert the ‘normal’ signaling in the brain environment to favor tumor growth. An excellent example of such a ‘hijack’ was recently discovered via a glioma-neuronal network that promotes gliomagenesis (Venkataramani et al., 2019; Venkatesh et al., 2019). Our findings suggest a role of MES-like cancer cells in ‘training’ infiltrating immune cells to acquire immunosuppressive traits (e.g., NECTIN, PROS pathways) and enhance neovascularization. The failure of anti-angiogenic therapies may be due to the monomodal targeting (anti-VEGF agents) of the tumor neoangiogenesis. Our results hint that the pro-angiogenic signaling may not solely depend on VEGF but on multiple redundant pathways (e.g., CALCR, ANGPTL). Skewing the cancer cells towards an MES-like phenotype (e.g., by employing OSM or functional analogs) (Hara et al., 2021), followed by targeted elimination, might decrease tumor-supporting activities of the immune cells and the pro-vascularization signaling. Successful hampering of the angiogenic signaling will require additional targeting of other components of the TME (e.g., TAMs, neutrophils) that further support this process.

Integrating single-cell and spatially resolved transcriptomics (Visium), we uncovered an unprecedented territorial organization of the GB TME. We could corroborate and extend the spatial disposition of the cancer cell states at single-cell resolution using HybISS. The exceptional ‘patches’ formed by MES-like malignant cells alongside a neighboring preference of AC- with OPC-like indicate possible environmental cues necessary to preserve its architecture. GB patterning appears intricately defined by landmark anatomical structures inside the tumor, in this case, the tumor vasculature. We observed an ‘anatomical’ allocation of the neoplastic states alongside blood vessels. Because of their considerable distance to the tumor vessel, the oxygen surrounding MES-like hubs is likely limited and may impact cell-cell communication. Hypoxia can activate signaling pathways that regulate epithelial to mesenchymal transition (Hapke and Haake, 2020). Whether hypoxia induces a ‘glial-to-mesenchymal’ shift or cancer cells’ fast growth and energy consumption cause hypoxia and subsequent upregulation of pro-angiogenic signaling, followed by microvascular proliferation, need to be addressed. Lastly, HybISS disclosed a propensity of the AC-like cells to surround the blood vessels, which could be either a feature of its astrocyte commitment (Hösli et al., 2022) or part of a blood-tumor barrier.

In agreement with Gangoso and colleagues, who recently showed how epigenetic changes in MES-like cells favor the recruitment of circulating myeloid cells (Gangoso et al., 2021), we found that the MES-like compartment co-localizes with monocytes/TAM-BDM. This preferred ‘partnership’ could result from tumor habitat conditions, as demonstrated recently by specific spatiotemporal patterns of GB TAMs driven by vascular alterations and tumor hypoxia (Sattiraju et al., 2022). The spatial co-occurrence of certain cancer states and immune populations is a feature of the TME that requires further investigation and could impact the development of new drugs, particularly immunotherapies. Spatial profiling of an increasing number of tissues will pave the way toward a better understanding of the GB dynamics and advance our knowledge for developing successful therapies with a future vision of personalized treatments.

The GBmap is a dynamic framework combining the single-cell transcriptomics of over 1 million cells that will benefit the scientific community by providing the opportunity to map new data, update the current model, scrutinize novel concepts, and generate hypotheses that can be explored experimentally.

## ACKNOWLEDGMENTS

C.R.M., E.S., M.N., S.B., and H.G.S. are supported by the European Union’s Horizon 2020 Skłodowska-Curie Actions (project AiPBAND) under grant #764281. C.R.M. and H.G.S. are additionally supported by the Princess Maxima Center and Kika (Kinderen Kankervrij Fast Track to H.G.S). S.B. is in addition supported by Edinburgh-UCL CRUK Glioma Centre of Excellence (CRUK-GCoE). M.N. also received funding from the Knut and Alice Wallenberg Foundation (KAW 2018.0172), the Erling Persson Foundation, the Chan Zuckerberg Initiative (SVCF 2017-173964), Cancerfonden (CAN 2018/604), and The Swedish research council (2019-01238). We thank A. Rios, P. Wesseling, and D. van Vuurden for their critical comments and suggestions on the manuscript. We thank K. Johnson and R.G.W. Verhaak, who shared single-cell RNA sequencing data before the formal publication of their study.

## AUTHOR CONTRIBUTIONS

C.R.M. and H.G.S. conceived, coordinated, and designed the study. C.R.M. generated single-nuclei RNA sequencing data from GB specimens. E.S. performed HybISS experiments. C.R.M. and S.M.S performed bioinformatic analysis and data visualization. C.R.M., E.S., and S.M.S analyzed data. C.R.M., M.N., and H.G.S. designed experiments. S.B. identified and consented patients for the study. M.K. assessed H&E staining of published ST Visium experiments. C.R.M. and H.G.S. wrote the manuscript, with feedback from all authors. All authors read and approved the final manuscript.

## SUPPLEMENTARY INFORMATION

Figures S1-S6

Tables S1-6

## METHODS

### RESOURCE AVAILABILITY

#### Lead Contact

Further information and resource requests should be directed to and will be fulfilled by the Lead Contact, Hendrik G. Stunnenberg.

#### Materials Availability

This study did not generate new unique reagents.

#### Data and Code Availability

The newly generated snRNA-seq data (processed and filtered) have been deposited in GEO under accession number GSE211376. The core and extended GBmap (raw and normalized counts, integrated embedding, cell type annotations, technical and clinical metadata) is publicly available and can be downloaded via cellxgene at https://cellxgene.cziscience.com/collections/999f2a15-3d7e-440b-96ae-2c806799c08c. Files for reference mapping (Azimuth), transfer learning (core reference model - scArches), and cell-cell interactions (CellChat) were made available at https://doi.org/10.5281/zenodo.6962901.

The published datasets that were included in the core GBmap can be accessed under GEO/EGA/SRA accession numbers: GSE103224, GSE131928, GSE138794 (EGAS00001002185, EGAS00001001900, and EGAS00001003845), GSE148842, GSE139448, EGAS00001004422, PRJNA579593, GSE117891, GSE157424, EGAS00001004656, EGAS00001005300, GSE84465, PRJNA588461, GSE135437, GSE163108, and GSE163120. Datasets included in the extended GBmap can be accessed under GEO/links: GSE141946, GSE166418, GSE162631, GSE154795, GSE141383, GSE182109, GSE173278, https://portal.gdc.cancer.gov/projects/CPTAC-3, and https://doi.org/10.17605/OSF.IO/4Q32E. Expression and metadata of the healthy brain dataset used for comparison with the GBmap are publicly available at http://stab.comp-sysbio.org.

Published spatially resolved transcriptomics datasets (Visium) analyzed in this study can be downloaded at Datadryad: https://doi.org/10.5061/dryad.h70rxwdmj. The expression maps and identity of each cell generated in this study using HybISS can be found at https://doi.org/10.5281/zenodo.6954130.

Details of the code and parameters used to create the core and extended GBmap, including downstream analyses (cell-cell interactions, spatial transcriptomics deconvolution), will be made available at https://github.com/ccruizm/GBmap.

### EXPERIMENTAL MODEL AND SUBJECT DETAILS

#### Human samples and tumor tissue sectioning

GB tissue sections from primary resections of adult patient tumors were obtained via Dr. Sebastian Brandner as part of the UK Brain Archive Information Network (BRAIN UK), funded by the Medical Research Council. Relevant ethical consent was provided by BRAIN UK (Ref:19/005). Fresh tumor tissue was collected and frozen immediately. After embedding in OCT, 20 µm sections were cut from the OCT blocks and mounted on positively charged slides (VWR superfrost Plus slides). The slides were then stored at -80° C until use.

### METHOD DETAILS

#### Single-nuclei RNA sequencing

Nuclei isolation was performed on 10-20µm cryosections for each specimen. The tissue slices were placed on 500µl of Nonidet P40 with salts and Tris (NST) lysis buffer1 and homogenized on ice using a glass-on-glass Dounce homogenizer with ten strokes using the loose pestle, followed by 15 strokes of the tight pestle and incubated for 5 min on ice. Nuclear homogenates were filtered through a 70µm Flowmi cell strainer (Bel-Art) and centrifuged for 5 min at 500g, at 4 °C. The pellet was resuspended in wash buffer1 and filtered with a 40µm Flowmi cell strainer (Bel-Art). Nuclei were stained with DAPI 1:200 dilution (2µg/ml stock concentration; Sigma-Aldrich) 5min at room temperature (RT) before sorting. FANS was done on a Sony SH800 cell sorter (Sony Biotechnology) using a 100µm nozzle. Single Cell Gene Expression 3’ v3 (10x Genomics) was used for single-cell/nuclei capturing and library construction, as described in the Genomics Single Cell RNA Reagent Kits User Guide. Briefly, 15,000 single sorted nuclei were loaded into a channel of a Chromium Single Cell Gene Expression 3’ Chip. Single nuclei were partitioned into droplets with gel beads in the Chromium, followed by barcoded reverse transcription of RNA, cDNA amplification, fragmentation, and sample index ligation. The quality of the libraries was assessed on a 2100 Bioanalyzer (Agilent) and sequenced on a NovaSeq (Illumina).

#### Gene Selection for HybISS

Gene panels were curated manually and computationally. The panels were based on snRNA-seq data from 11 GB patients. Gene selection was first established on differential expression between respective cell types, followed by manual filtering of genes with likely high background expression levels being strongly expressed in all cell types and favoring genes established as cell type markers in classical immunohistochemistry. A total of 954 individual probes were designed from a selection of 194 genes encompassing malignant (OPC-like, AC-like, NPC-like, and MES-like) and non-malignant cells (microglia, macrophages, oligodendrocytes, astrocytes, neurons, endothelial cells, T cells, dendritic cells, and mural cells), as well as signaling markers of interest. Malignant cell type definitions were defined using snRNA-seq datasets via copy-number variation (CNV) inference for cells that saw chromosome 7 gain and 10 loss. In contrast, non-malignant cell type definitions were defined from the same dataset with no chromosomal aberrations.

#### Probe Design for HybISS

Padlock probes were designed for the selected genes, each containing two arms matching a 40- base-pair (bp) sequence on the cDNA, a 4-bp barcode, an ’anchor sequence’ allowing all amplicons to be labeled simultaneously, and a 20-bp hybridization sequence for additional readouts. Target sequences for the selected genes were obtained using in house Python padlock design software package that utilizes ClustalW and BLAST+ (https://github.com/Moldia/multi_padlock_design) with these parameters: arm length, 15; Tm, low 65, high 75; space between targets, 15. After target sequences were obtained, five targets were selected randomly per gene. If fewer targets were found, then only those were selected. The backbone of the padlock probes (PLPs) includes a 20 nucleotide (nt) ID sequence and a 20 nt sequence ’anchor’ that is common among subsets of PLPs (Note: In this study, the anchor sequences were not used and served only as linker sequences.)

#### Hybridization-based In Situ Sequencing (HybISS)

HybISS was performed as described previously(Gyllborg et al., 2020). Briefly, after fixation with 3% PFA for 30 min, the sections were then permeabilized with 0.1 M HCl and washed with PBS. After being rehydrated for 1 minute in 100% ethanol, 1 minute in 75% ethanol, and 1 minute in PBS. cDNA synthesis was run via reverse transcription overnight with reverse transcriptase (BLIRT), RNase inhibitor, and priming with random decamers. Sections were then post-fixed before PLP hybridization and ligation at a final concentration of 10 nM/PLP., with Tth Ligase and RNaseH (BLIRT). This was run at 37°C for 30 minutes and then moved to 45°C for 1 hour. Sections were washed with PBS, and rolling circle amplification (RCA) was performed with phi29 polymerase (Monserate) and Exonuclease I (Thermo Scientific) overnight at 30°C. The designed bridge probes (10 nM) were hybridized at RT for 1 h in the hybridization buffer (2XSSC, 20% formamide), followed by the hybridization of readout detection probes (100 nM) and DAPI (Biotium) in the hybridization buffer for 1h at RTThe sections were washed with PBS and mounted with SlowFade Gold Antifade Mountant (Thermo Fisher Scientific). After each imaging cycle, the coverslips were removed, and sections were washed 5 times with 2XSSC, and then bridge probe/detection oligonucleotides were stripped with 65% formamide and 2XSSC for 60 min at 30°C. This was followed by 5 washes with 2XSSC. Then the above procedure was repeated for cycles 1 through 5, with each cycle consisting of cycle-specific individual bridge probes to be hybridized as above.

#### Imaging

Imaging was performed using a standard epifluorescence microscope (Zeiss Axio Imager.Z2) connected to an external LED source (Lumencor® SPECTRA X light engine). The light engine was set up with filter paddles (395/25, 438/29, 470/24, 555/28, 635/22, 730/40). Images were obtained with an sCMOS camera (2048 × 2048, 16-bit, ORCAFlash4.0 LT Plus, Hamamatsu), automatic multi-slide stage (PILine, M-686K011), and Zeiss Plan-Apochromat objectives 20x (0.8 NA, air, 420650-9901), 40× (1.4 NA, oil, 420762–9900). Filter cubes for wavelength separation included quad band Chroma 89402 (DAPI, Cy3, Cy5), quad-band Chroma 89403 (Atto425, TexasRed, AlexaFluor750), and single band Zeiss 38HE (AlexaFluor488). Each field-of-view (FOV) was imaged with 21 z-stack planes with 0.5 μm spacing and 10% overlap between FOVs.

### QUANTIFICATION AND STATISTICAL ANALYSIS

#### Processing of published data and construction of the core GBmap reference

For the construction of the core GBmap, only samples confirmed to be GB IDH wild-type (based on the clinical metadata provided in each study) and containing at least 1000 cells were included. Transcriptomic data coming from nuclei were not considered for the training of the reference model. The 16 datasets collected in the core GBmap (Table S1) were obtained either as raw or TPM normalized (for Smart-seq2 studies) count matrices. In studies where raw count matrices were unavailable (Couturier2020, Bhaduri2020), BAM files were converted to FASTQ files and re-aligned to GRCh38 using CellRanger v3.1/4.0. All gene names were updated to the latest HUGO nomenclature using HGNChelper (Oh et al., 2020). All clinical/diagnostic metadata was harmonized and preserved.

Before integrating the datasets, we applied homogeneous filtering parameters to include high-quality cells, excluding cells that expressed fewer than 500 genes, 1000 UMI counts (for datasets where applicable), and more than 30% mitochondrial reads. We estimated and discarded potential doublets for each droplet-based dataset using DoubletFinder (McGinnis et al., 2019). To reduce the batch effects among datasets, we used a semi-supervised neural network model of probabilistic harmonization denominated single-cell ANnotation using Variational Inference (Xu et al., 2021) (scANVI) under the transfer-learning model implemented in the single-cell architectural surgery algorithm (Lotfollahi et al., 2021) (scArches). As a semi-supervised model, scArches-SCANVI requires prior knowledge of the cell types/labels when creating the reference map. To harmonize cell type labels from different sources, we annotated each dataset using automated and manual methods (Clarke et al., 2021). For the automatic cell annotation, we compiled a curated gene marker list from different studies (Table S6), providing a signature for cell types part of the GB TME and used it as input for the Cell-ID algorithm (Cortal et al., 2021). After, for the manual assignation of cell identity, we considered the results from the automatic cell annotation, the original cell label (when available), and cell-type annotation available on the TISCH website (Sun et al., 2021) (when available), and the expression of cell type-specific marker genes identified using the Wilcoxon rank-sum test by comparing all cells within a specific cluster to all cells outside said cluster. Particularly for the accurate identification and annotation of neoplastic cells, for all datasets but immune enriched, CNV inference was carried out using the CopyKAT package (Gao et al., 2021), classifying cells that were either diploid or aneuploid. This preliminary coarse cell type labeling was used for the model training and integration via scANVI-scArches. The pipeline was run on the raw counts of the 5000 most highly variable genes (HVGs), using studies as the batch variable and the recommended parameters of the tool. The pipeline output is the latent representation of the integrated data that serves as input for clustering and dimensional reduction visualization. We used a k-nearest neighbor graph (k-NNG)-based Leiden clustering (Traag et al., 2019) to detect the distinct cell populations and Uniform Manifold Approximation and Projection (UMAP) (Becht et al., 2019) for data embedding and two-dimensional reduction. UMAP visualization of the core GBmap (**Figure 1b**) was generated using the plot1cell package (Wu et al., 2022). Total count normalization was done by initially dividing each count by the total count per cell and multiplying by 10,000, followed by log transformation using a natural *log*(*X* +1).

After co-embedding all cells, we refined the cell annotation by manually assigning a cell identity to each cluster, considering our unified preliminary cell annotation and the expression of specific marker genes that correctly defined each broad cell type/state (level-1, -2, and -3 annotation). To determine the next level of cell identity (level-4 annotation), we sub-selected, re-clustered, and identified the primary axes of transcriptional variation on each cell territory of interest (malignant, lymphoid, myeloid, and vascular) using Hotspot (DeTomaso and Yosef, 2021), which allowed the identification of genes that vary in a contextualized fashion. The output is the organization of the genes of each territory into co-varying groups (module/program). We ran the tool using the latent space as inferred by scVI (Lopez et al., 2018) to build the k-NNG and used it as input of the source of cell-cell distances to compute the Euclidean distance in the low-scVI-dimensional space. Hotspot employs a negative binomial distribution to compute pair-wise local correlations between the top500 lineage autocorrelated genes and group them into correlated modules. Each gene module was matched with a thorough literature search of specific phenotypes that delineated a given cell state. These included findings made by previous studies in high-grade gliomas or extended to other cancer types if there was no match in other brain-related pathologies. After calculating the enrichment of each cell for every module, the phenotype assignment was performed based on the highest score of a cell for a given gene program.

#### Data processing of *de novo* GB samples

Raw BCL files generated by the sequencer were demultiplexed using Cell Ranger mkfastq (v3.1.0) to generate the FASTQ files. Each sample was mapped to the human reference genome (GRCh38 v3.0.0) provided by 10x Genomics using the Cell Ranger count with default parameters to obtain the gene count matrix. For single nuclei samples, the reference for pre-mRNA was created using the manufacturer’s guidelines (https://support.10xgenomics.com/single-cell-gene-expression/software/pipelines/latest/advanced/references).

#### Query projection onto the GBmap core

For the projection and annotation of external data sets onto the GBmap, we first created an object that could be manipulated and utilized by the Azimuth algorithm part of the Seurat package (Hao et al., 2021). We imported the Anndata object into R and created a Seurat object that contained the raw and normalized count matrix, cell metadata, and UMAP embedding. The Azimuth label-transferring method uses the low-dimensional structure of the reference and finds ’anchor’ genes on the query dataset that enables the projection and transfer of the cell annotations between data sets. For the eleven patients profiled by snRNA-seq in our study, we used unsupervised anchoring based on the first 50 principal components and the log normalized matrices. Each cell in the query dataset gets a prediction probability, and the cell assignment is performed based on the highest score for a given cell type/state. To confirm the correct assignment of neoplastic and non-neoplastic cells, we used the inferCNV package (Tickle et al., 2019) placing as reference cells that were not expected to carry any CNV, such as immune and vascular cells. Detection of marker genes among predicted cell types and states was done using a Wilcoxon rank-sum test.

#### Expansion of the GBmap by transfer learning

To extend the GBmap, we gathered raw count matrices from newly generated public datasets available after the curation of the core reference atlas (April 2021) (Table S1). By using the deep learning scArches algorithm (Lotfollahi et al., 2021), the core GBmap cannot only allow re-annotation of queried cells but also update the currently trained model and capture differences that could represent new cell types or states. In short, scArches trains adaptors that are added to the reference embedding model, which facilitates the creation of a de novo joint embedding between the new datasets and the core GBmap, enabling the generation of a new dimensional reduction and clustering and re-analysis of the extended reference. The new datasets were QC filtered using the same parameters previously established for constructing the core GBmap. We merged the data matrices of the different studies and selected the same 5000 HVGs employed for the training of the reference model. Raw counts were used as input for scArches. The integration pipeline was run to ’re-train’ adapter weights, thus enabling the mapping of new query data into the core GBmap dimensional embedding. To find DEGs between cell types, we used a Wilcoxon rank-sum test comparing cells in the core reference with a specific cluster from the updated version of the GBmap.

#### Cell-cell interaction analysis in the GBmap

To infer the cell-cell communication among the cell types in the GB TME, we utilized the CellChat package (Jin et al., 2021). We created a custom L-R database (DB) by merging the ’CellChatDB.human’ L-R interaction default DB and FANTOM5 resource (Ramilowski et al., 2015), increasing the amount of curated L-R interactions and grouping them in meaningful biological pathways, broadly divided into ’Secreted Signaling’, ’ ECM-Receptor’ and ’Cell-Cell Contact’ interactions. The custom DB was used to compute the communication probabilities among the cell types in the healthy brain and extended GBmap (level-3 annotation) datasets individually. Intercellular communication networks are weighted directed graphs composed of significant interactions among cell populations, where the interaction strength is defined as the communication probability of the computed communication pathways. Differential interactions between both conditions were calculated following the recommended comparison analysis of multiple datasets with different cell type compositions. Due to the vastly distinct cell composition between healthy and brain tumors, pair-wise similarity analysis can be done only by assessing the differences and changes in the structural network topology. The information flow displays the overall communication probability results of the probability among all pairs of cell groups in the inferred network, and pathway distances were determined by computing the Euclidean distance between the pairs of the shared signaling networks across the two conditions (normal brain vs. GB). We provide a CellChat object of the GBmap interactome that can be explored interactively using the CellChat R package.

#### Pathway activity and gene set enrichment analysis (GSEA)

Inference of pathway activity was performed with PROGENy, using the implementation to analyze single-cell RNA-seq data (Holland et al., 2020). PROGENy infers pathway activity for 14 signaling pathways (Androgen, Estrogen, EGFR, Hypoxia, JAK-STAT, MAPK, NFkB, PI3K, p53, TGFb, TNFa, Trail, VEGF, and WNT) on matrices of single cells containing normalized gene expression values. By default, pathway activity inference is based on gene sets comprising responsive genes for a specific pathway. The result is a pathway score based on the strength of pathway regulation and gene expression. GSEA was performed using the R package fgsea (Korotkevich et al., 2021) (MsigDB hallmark gene set collection) on the gene expression signatures of each cluster resulting from the Wilcoxon test.

#### Deconvolution of cell types and analyses of Visium data

We downloaded from the 10x Genomics website a publicly available GB sample profiled using the Visium ST platform (https://www.10xgenomics.com/resources/datasets/human-glioblastoma-whole-transcriptome-analysis-1-standard-1-2-0), as well as nine high-quality GB tissues profiled on the Visium platform (Ravi et al., 2022). Data importing, basic filtering of spots based on total counts and expressed genes, normalization, dimensional reduction, clustering, and detection of spatially variable features were performed in Python using the Scanpy package (Wolf et al., 2018). Histopathological evaluation of the H&E staining to delineate key anatomical features on the tumor sections were made by M.K.

To spatially map cell types defined in the GBmap and determine the cell abundances in each capture area of the Visium ST data, we used a Bayesian model implemented in the cell2location tool (Kleshchevnikov et al., 2022) that decomposes coarse ST data into cell-type abundance estimates in a spatially resolved manner. cell2location derives expression signatures of cell types in the scRNA-seq reference. In the case of the GBmap, we extracted the expression signatures on the extended GBmap (level-3 and -4 annotation) using the negative binomial regression model implemented in the tool that allows a robust integration of expression profiles across technologies and batches. To obtain cell-type distributions, cell2location performs a non-negative decomposition of the gene signatures in every spot and matches the cell type profiles defined in the reference. Each ST section was analyzed independently. Cell2location parameters were set to default values. The spatial maps show the estimated cell abundance (5% quantile, representing confident assignment to each cell type) of each GBmap population in each spot.

To define spatial compartments, we applied a non-negative matrix factorization (NMF) of the absolute cell type abundance estimates per spot obtained from cell2location, considering all analyzed Visium samples. To estimate diversity in each capture location on the Visium slides, we employed the stLearn package (Pham et al., 2020), which takes the estimated cell abundances derived from cell2location to calculate the cell mixture of each spot. Spatial L-R interaction analysis was carried out through the Squidpy tool (Palla et al., 2022) and the NICHES package (Raredon et al., 2022), employing the extended and curated Omnipath L-R DB (Ceccarelli et al., 2020).

#### Image processing and decoding of HybISS data

Each FOV image was maximum intensity projected to obtain flattened two-dimensional images. These images were then analyzed with in-house custom software. Each two-dimensional FOV was exported, aligned between cycles, and stitched together using the MIST algorithm. Stitching was followed by retiling to create smaller non-overlapping 6000x6000 pixel images that were then used for decoding. The decoding pipeline can be found on the Moldia GitHub page (https://github.com/Moldia/iss_starfish/). Using Starfish, images were initially filtered by applying a white top hat filter. The filtered images were subsequently normalized, and spots were then detected using the FindSpots module from Starfish, which were then decoded using the MetricDistance module. Finally, the resulting spots were filtered based on the distance to the closest expected barcode in the reference codebook.

#### Cell typing and spatial statistical analysis of HybISS

Probabilistic Cell Typing (pciSeq) (Qian et al., 2020) was used to identify the cellular identity of both malignant and nonmalignant cells based on the prior expression signatures of each of the cell types detected on each sample. Using this approach, each cell was given a probability of belonging to each pre-defined cell type, including a signature for cells with too low counts, which were then excluded from the analysis. Spatial statistics, including neighborhood enrichment, were generated using Squidpy (Palla et al., 2022). In short, the neighborhood enrichment between cell type A and B represents the probability of assigning a cell-to-cell type A given the presence of a second cell of cell type B within a certain distance (*d)* divided by the probability of assigning a random cell to cell type A. To explore the enrichment of specific cell types close to areas with cells expressing genes of interest (*VEGFA, HIPLDA)*, the same algorithm was applied, computing the neighborhood enrichment of particular cell types and gene reads with a certain identity. In order to represent the effect of *d*, we represented the changes in neighborhood enrichment scores of certain pairs (y-axis) when modifying the distance (x-axis). To explore the cellular environment of each cell and define cellular niches, each cell was re-defined based on the identity of the cells situated within 50um close to itself, creating a Cell-by-neighboring cell types matrix. After normalizing and log-transforming the counts, UMAP and Leiden clustering were performed to identify cellular niches.

#### Statistical analysis

The statistical analyses were performed using R, Python, and MatLab.

## SUPPLEMENTAL INFORMATION

### Supplemental Figure Legends

**Supplemental Figure 1.**
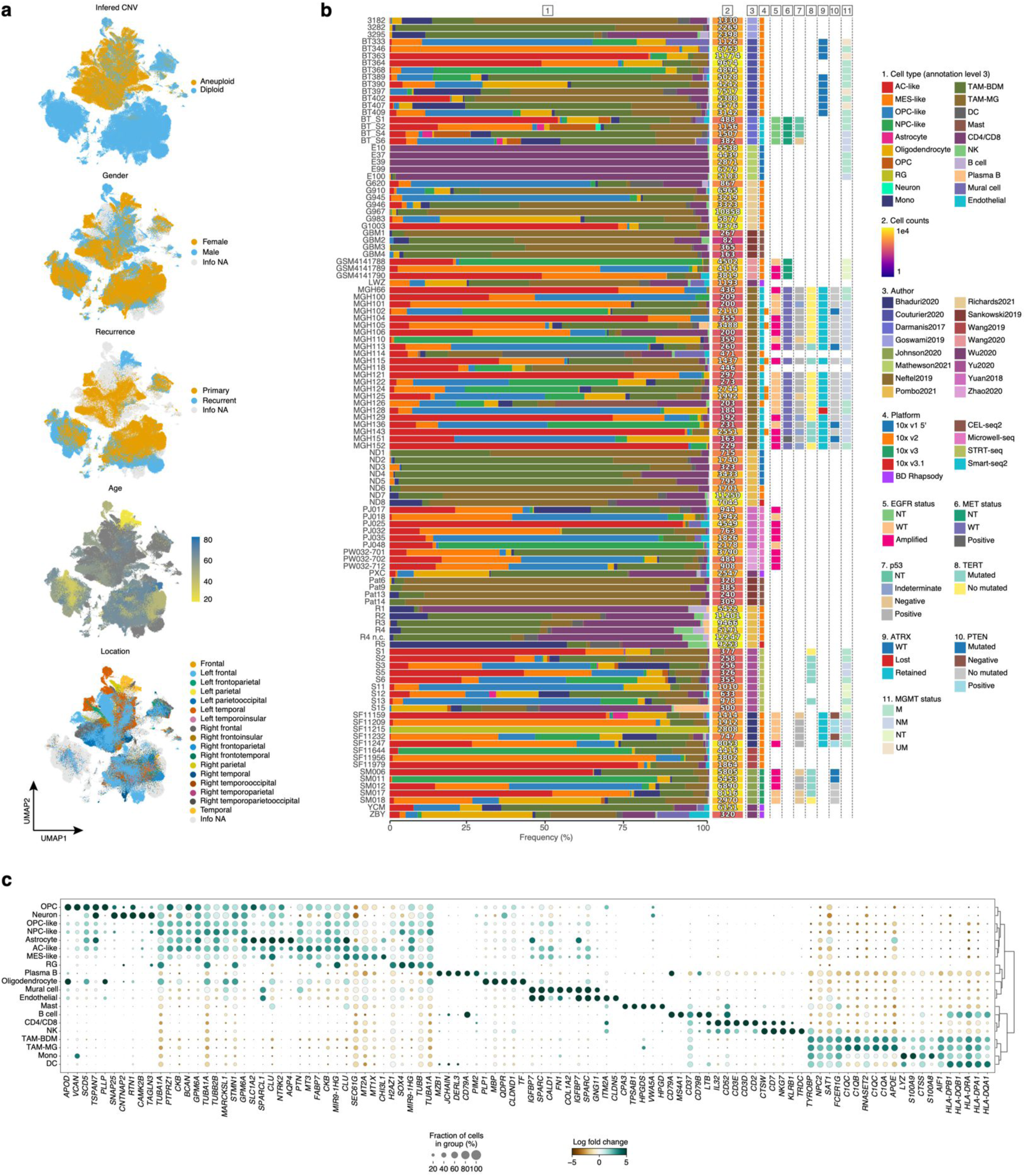
Construction of the GB TME landscape, related to Figure 1. (a) UMAP of the GB atlas colored by iCNV and clinical features. (b) Bar plot of cell distribution, cell count, study, platform, and genomic features reported per patient. (c) Gene expression of top5 genes from the level-3 annotation resolved on the GBmap.

**Supplemental Figure 2.**
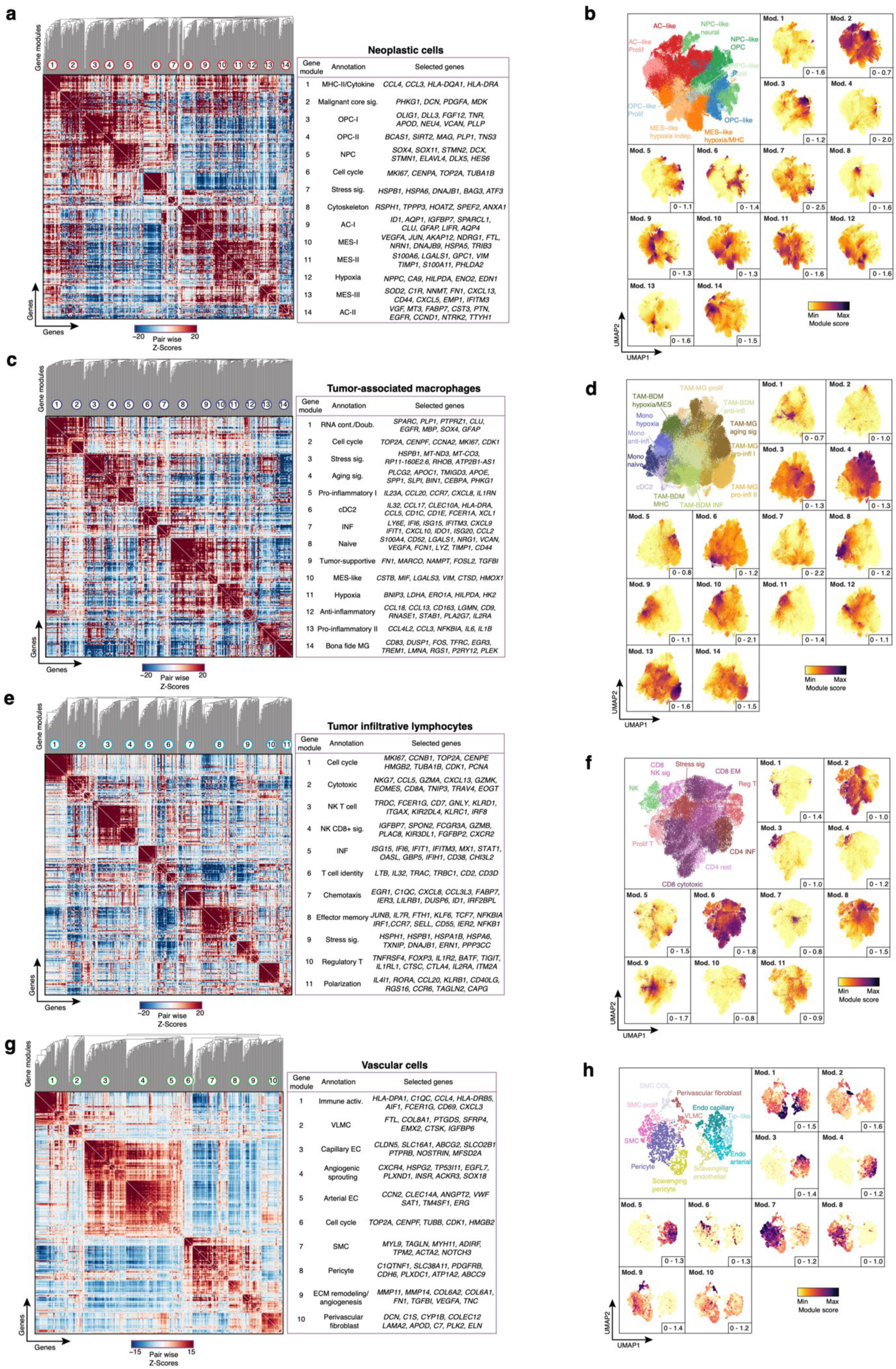
Definition of cell states in the GB TME, related to Figure 2. (a, c, e, and g) Heatmap of gene pairwise local correlation of the top 500 genes detected in neoplastic cells, TAMs, TILs, and vascular cells, grouped into gene modules. (b, d, f, and h) UMAP of sub-clustered neoplastic cells, TAMs, TILs, and vascular cells, and gene module’s enrichment score. The minimum and maximum scores of each module are shown in the bottom right of each panel.

**Supplemental Figure 3.**
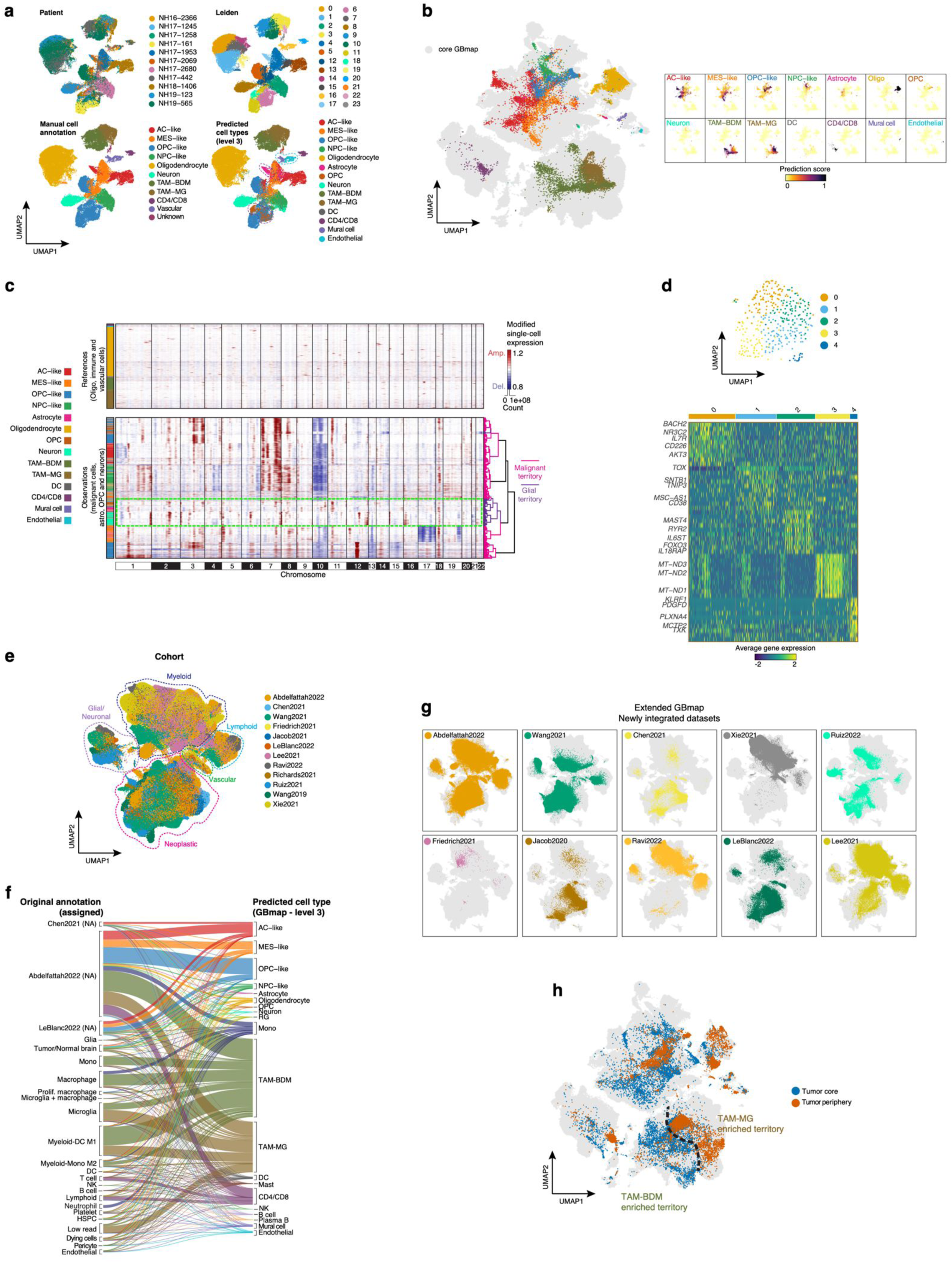
Projection and integration of query datasets onto the GB atlas, related to Figure 3. (a) UMAP is colored by patient, Leiden clustering, manual, and predicted cell annotation from the GBmap (11 samples processed in this study), using an anchoring method for reference mapping (Azimuth). (b) Projection of cell types (level-3 annotation) of *de novo* mapped query cells (11 samples processed in this study) onto the core GBmap UMAP and respective prediction scores. (c) Heatmap of iCNV profiles of the query dataset (this study). Color bar shows the predicted cell type signature for each cell. Green dotted line emphasizes neuronal/glial cells lacking iCNV. (d) UMAP of reclustered TILs from *de novo* GB dataset (11 samples processed in this study) and top gene expression for each subcluster. (e) UMAP of integrated public datasets and our own cohort. Colored by cohort and dotted lines enclose the main cell territories. (f) Comparison of original annotation with predicted cell types (level-3 annotation) after label transfer from the GBmap core. (g) UMAP of the extended GBmap colored and split by queried studies. (h) UMAP displays the distinction between core and periphery regions of multi-sector biopsy studies included in the GBmap. The dashed line indicates the estimated boundary between the TAM-BDM and TAM-MG territories.

**Supplemental Figure 4.**
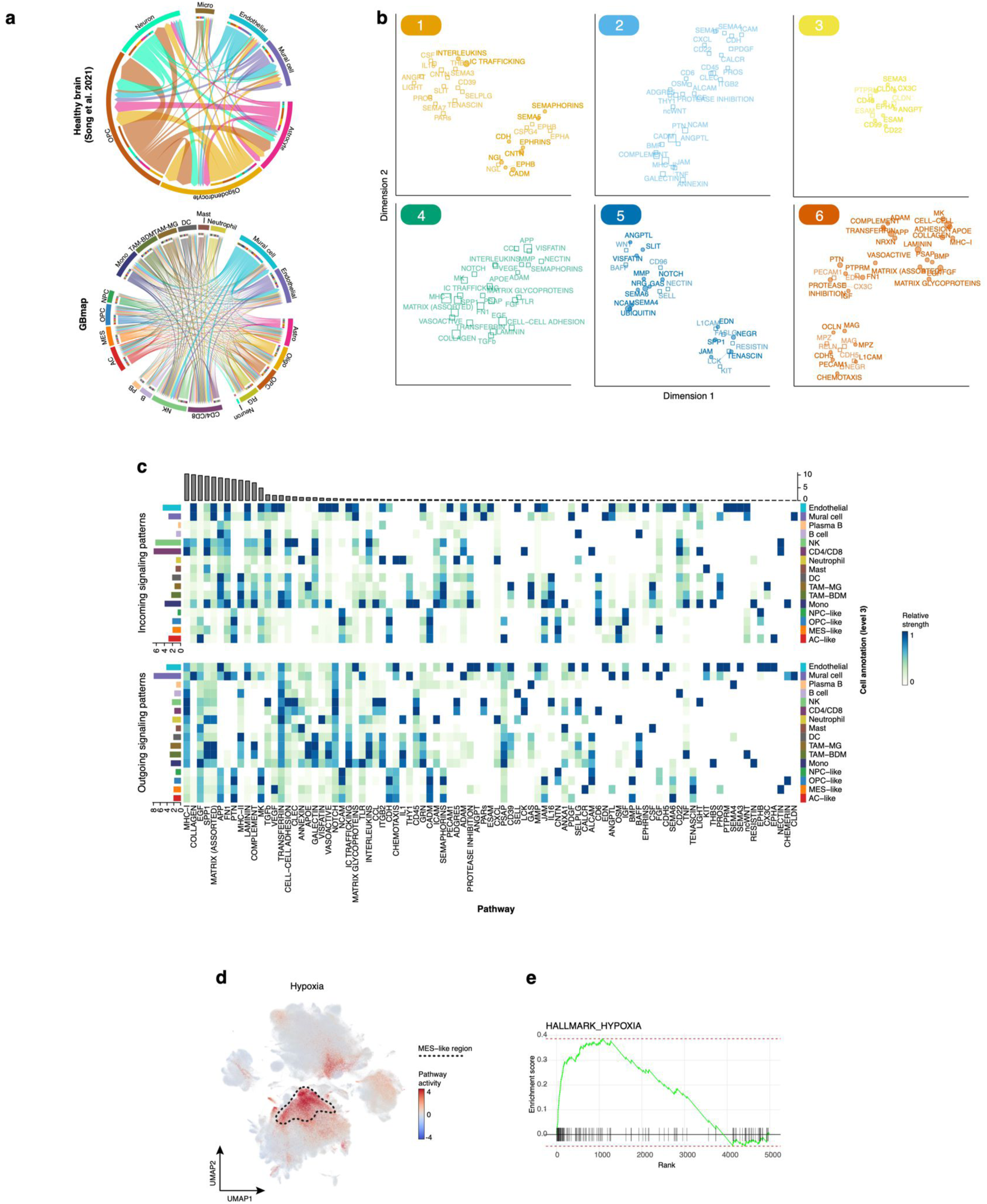
Cell-cell interaction network in GB, related to Figure 4. (a) Chord diagrams of statistically significant pathways (ligand-receptor) interactions among cell types in healthy brain and GB. (b) Magnified view of each pathway group from the shared two-dimensional manifold of healthy and tumor brain tissue signaling networks. (c) Heatmap of aggregated outgoing or incoming signaling of the cell-cell communication network from the GB signaling pathways. Bars display relative contribution per cell type and pathway. (d) UMAP of pathway activity prediction by PROGENY in the extended GBmap. The dashed line highlights the MES-like malignant area. (e) GSEA analysis revealing a hypoxia enrichment score for MES-like malignant cells.

**Supplemental Figure 5.**
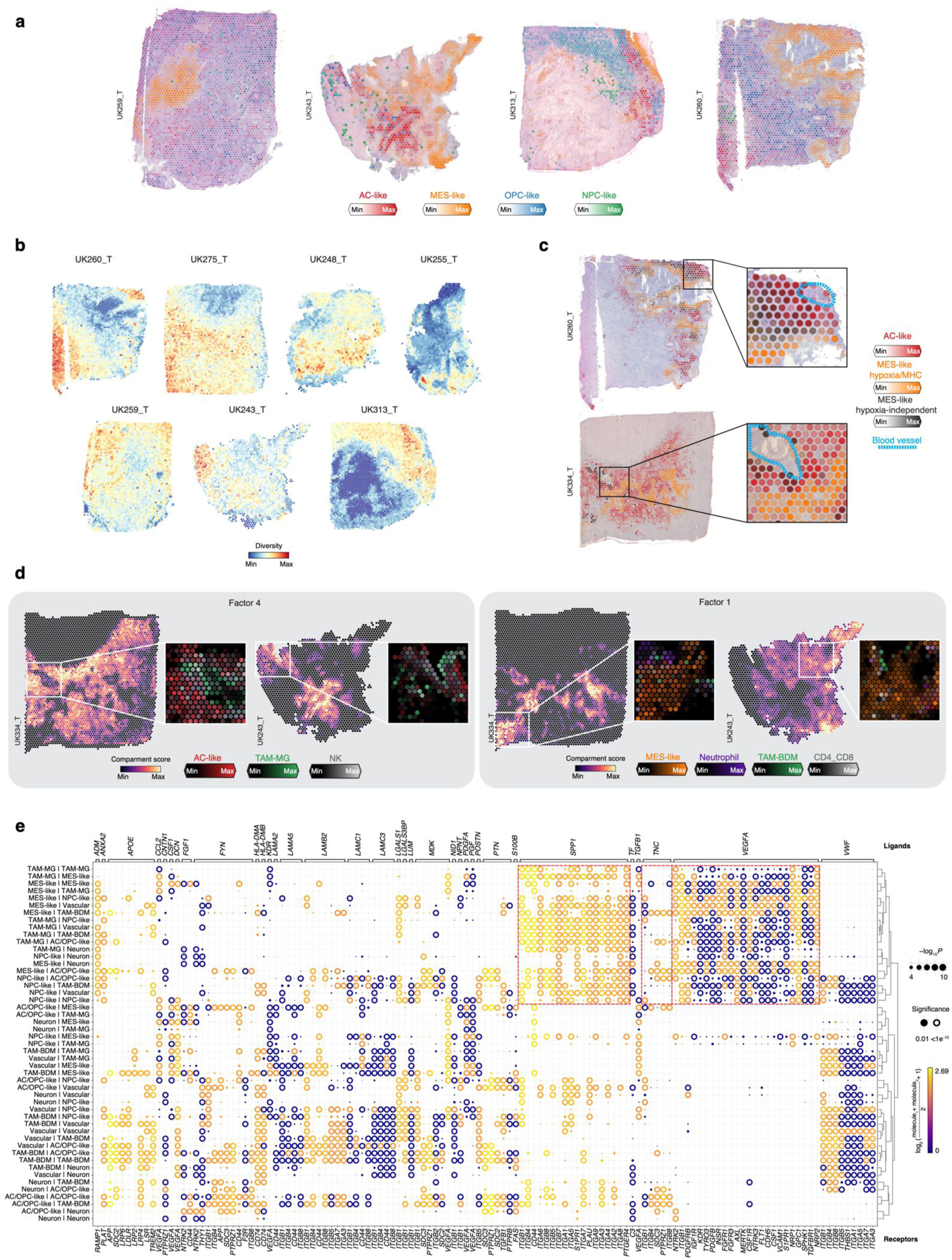
Spatial deconvolution and intercellular communication of NGS-based GB datasets, related to Figure 5. (a) Estimated cell abundance per spot (color intensity) of the four main cancer cell states shown over the H&E image of four additional primary GB Visium samples. (b) Cell type diversity within spots based on inferred cell type abundances per location. (c) Estimated cell-type proportions (color intensity) of selected cancer subtypes on two GB. Boxes highlight the areas shown in the inset panels. Dashed blue line depicts localization of the tumor vasculature. (d) Spatial plots depicting enrichment of the NMF compartments (factor 1 and 4; color intensity) in two GB samples. Boxes highlight the areas shown in inset panels and are colored by the estimated cell abundance (color intensity) of selected cancer states and immune cells. (e) Inferred R-L interactions from gene expression among defined spatial cell types, clustered by interacting clusters on NGS-based datasets.’

**Supplemental Figure 6.**
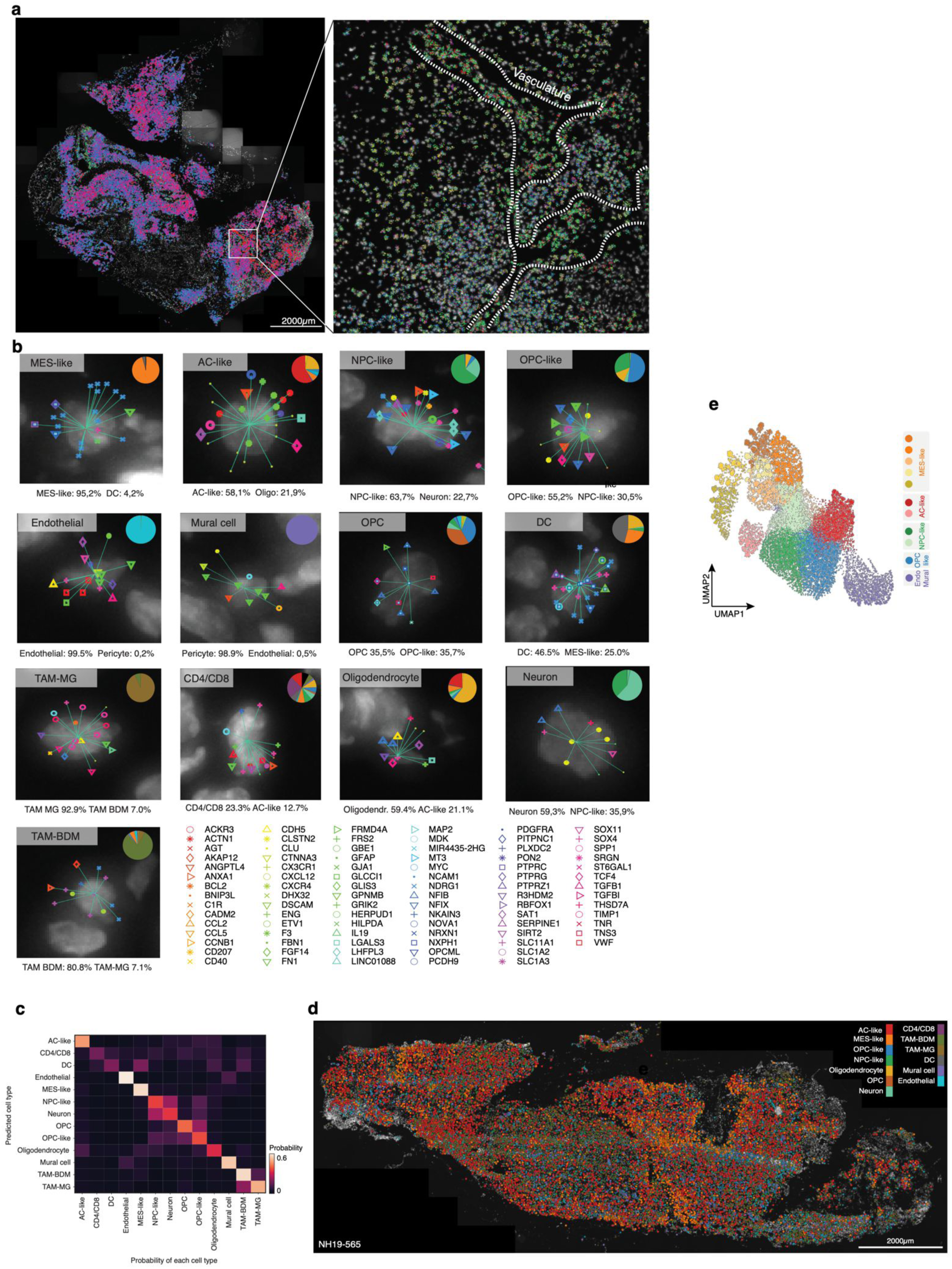
pciSeq of GB samples, related to Figure 6. (a) Spatial map of individual gene signals across sample NH17-2680. Genes are represented by designated colors and shapes. The dotted line in the right-hand image represents a structure of interest denoted by vascularized areas consisting of pericytes and endothelial cells. (b) Reads are assigned to cells and cells to classes using a pciSeq. Images show the distribution and assignment of reads for thirteen example cells. Colored symbols indicate reads per each assessed gene. Grayscale background image indicates DAPI stain. Straight lines join reads to the cell, which are assigned based on the highest probability. The pie charts show the probability distribution of each class. Colors indicate broad cell types; segments show probabilities for individual scRNA-seq clusters (with the two most dominant named underneath each image). (c) Confusion matrix of pciSeq of cell type scores for GB biopsy section (NH17-2680). Colors represent the mean probability assigned to a cell when a specific cell type is predicted. (d) pciSeq analysis of GB tissue (NH19-565), outlining the distribution of cell types and states, colored by cell identity. (e) UMAP of spatial analysis of the composition of the neighboring cells identified within the range of 50 μm (see Methods), colored by sub-niches.

### Supplemental Tables

**Supplemental Table 1:** Features of datasets included in the core and extended GBmap.

**Supplemental Table 2:** Gene signatures employed to perform automated cell typing using CellID.

**Supplemental Table 3:** Gene modules of each main cell territory obtained with HotSpot.

**Supplemental Table 4:** Differentially expressed genes for each annotation level (1 to 4) of the core GBmap.

**Supplemental Table 5:** Patient metadata of newly profiled GB.

**Supplemental Table 6:** Sequences of the padlock probes (PLPs) designed for *in situ* sequencing experiments.

